# Selection of a *de novo* gene that can promote survival of *E. coli* by modulating protein homeostasis pathways

**DOI:** 10.1101/2023.02.07.527531

**Authors:** Idan Frumkin, Michael T. Laub

## Abstract

Cells sometime adapt to challenging environments by turning non-functional loci into functional genes in a process termed *de novo* gene birth. But how proteins with random amino acid sequences integrate into existing cellular pathways to provide a benefit remains poorly understood. Here, we screened ∼10^8^ random genes for their ability to rescue growth arrest of *E. coli* cells producing the ribonuclease toxin MazF. Approximately 2,000 random genes could promote growth by reducing transcription from the promoter driving *mazF* expression. Additionally, one gene, named random <u>a</u>ntitoxin of <u>M</u>az<u>F</u> (*ramF*), whose protein product was well-tolerated in cells and neutralized MazF by interacting with chaperones, leading to MazF proteolysis. We show that the specificity of RamF for MazF relative to other toxins relies on the degron-like function of MazF’s first 10 amino acids. Finally, we demonstrate that random proteins can improve during evolution by identifying beneficial mutations that turned RamF into a more efficient inhibitor. Our work provides a mechanistic basis for how *de novo* gene birth can produce new, functional proteins that are integrated into complex cellular systems and provide a benefit to cells.

## Introduction

A central premise in molecular evolution is that organisms must innovate to survive changing environments. Cellular novelty usually emerges via mutations to existing genes or by mixing-and-matching domains from different genes^1^. However, evolution may also invent novel functional proteins from scratch, a process termed *de novo* gene birth^2, 3^. Yet, little is known about how often this process occurs and, when it does, how such new proteins function and provide a benefit to cells^4^.

Recent studies have used comparative genomics and synteny-based methods to identify lineage-specific genes that may represent *de novo* genes^5–7^. However, the designation of lineage-specific genes as *de novo* genes suffers from high false discovery rates due to homology detection failure^8^ and the intrinsic shortcomings of genome annotation methods^9^. For *bona fide* cases of *de novo* genes, some biological effects have been reported^10^, but whether they have beneficial functions that confer a selective advantage remains unknown in most cases.

How can a random nucleotide sequence become a gene? The “proto-gene” model for *de novo* gene birth^11^ sets two main requirements: (i) stable expression of a locus, and (ii) beneficial function of the emerging gene product. If these two conditions are met, natural selection can further improve expression, function, and regulation to generate a mature gene that is integrated into cellular physiology. RNA-sequencing and ribosome profiling studies have revealed extensive spurious transcription and translation in species across the tree of life^7, 11–15^. These expressed loci could serve as a source for novel genes.

A complementary approach to investigating *de novo* gene birth involves the characterization of randomly generated proteins and studying whether they can benefit cells. Prior work has examined *in silico* and *in vitro* properties of such random sequences, including their predicted ability to fold into secondary structures^16^, solubility^17^, ATPase activity^18^, or potential affinity for different molecules^19–22^. However, cellular functions for random genes have rarely been demonstrated *in vivo*. Two recent studies identified a few random small proteins containing transmembrane domains that confer antibiotic resistance, likely by modulating membrane physiology^23, 24^. Perhaps as expected for hydrophobic proteins, expression of these random proteins was not well tolerated, leading to a substantial fitness cost.

The functions that random proteins, especially cytosolic ones, can assume inside cells remain poorly understood. To probe this question, we created and screened a library of ∼10^8^ random genes for their ability to promote cellular growth in the presence of the ribonuclease toxin MazF. We found ∼2,000 unique genes that restore growth to cells producing MazF. Although most function nonspecifically to affect the promoter driving *mazF* to reduce its transcription, we found a single Random Antitoxin of MazF, now called RamF, that specifically rescued cells from MazF toxicity. We characterized RamF’s function, specificity for MazF, and mutational pathways to becoming a more efficient inhibitor. Our experiments indicate that RamF is a well-tolerated cytosolic protein that remodels the physiology of *E. coli* cells by interacting directly with cellular chaperones, leading to MazF proteolysis. Thus, our work demonstrates how a small, random protein can instantly have a beneficial function, integrate with complex, pre-existing cellular pathways, and become improved by mutation and selection - thereby revealing a plausible mechanism for *de novo* gene birth.

## Results

### Selection for functional, random genes that inhibit the toxin MazF

We sought to identify genes originating from random nucleotide sequences that are functional and beneficial. To this end, we created a library of ∼10^8^ plasmids, each harboring a tetracycline-inducible promoter (P*_tet_*) driving a bi-cistronic operon with a first ORF encoding a constant 17-amino acid peptide followed by a second ORF with an ATG start codon and then 50 random NNB codons (Fig. 1A; see Methods). This bi-cistronic design minimizes translation initiation biases due to mRNA structures involving the second ORF^25^. Deep sequencing of the initial library demonstrated its high complexity, with 99.42% of the ∼370,000 reads being single, unique sequences (Fig. S1). The average length of the random open reading frames (ORFs) was 28 amino acids, with 23% of the random genes coding for 51-amino acid proteins (Fig. 1B).

**Figure 1.**
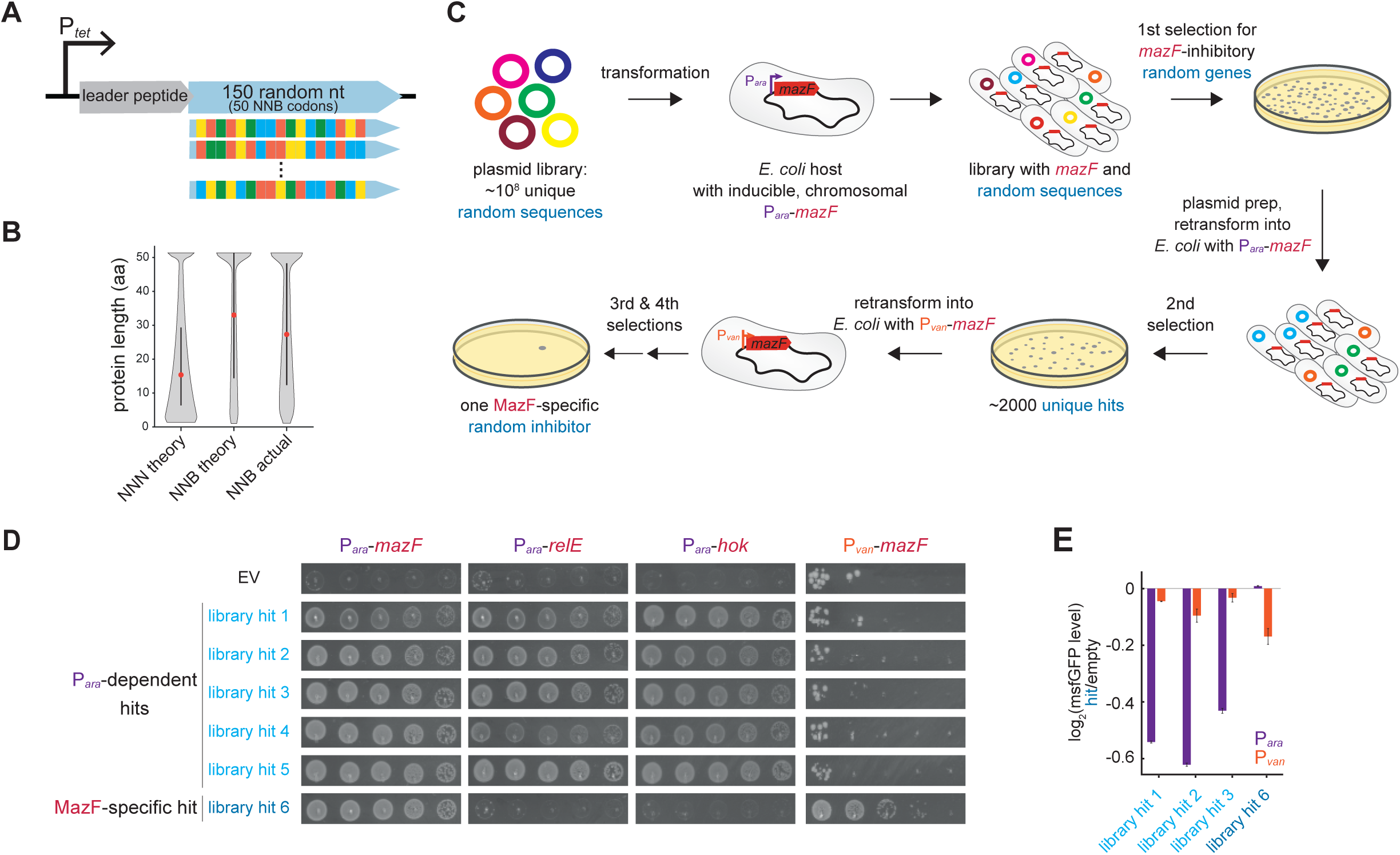
Strategy for selecting functional proteins from a random sequence library. (A) Architecture of the random sequence library. A tetracycline-inducible promoter (P*_tet_*) drives the expression of a leader peptide followed by an ATG start codon, 150 random nucleotides (50 NNB codons), a stop codon, and a transcriptional terminator. (B) Theoretical protein lengths of 50 NNN codons are lower compared to 50 NNB codons. Actual library distribution as deduced by deep-sequencing pre-selection is similar to the NNB distribution. Black bar represents 50% of variants and red dot is the median. (C) Selection strategy to identify functional proteins that inhibit the toxin MazF. Approximately 10^8^ plasmids harboring unique, random genes were transformed into an *E. coli* strain with a chromosomal, arabinose-inducible P*_ara_*-*mazF* gene. Surviving colonies after *mazF* induction include true hits and false positives due to chromosomal mutations. Plasmids were purified, re-transformed into new cells, and selected for a second time. The surviving colonies were then screened twice in a strain expressing *mazF* from the vanillate-inducible promoter (P*_van_*), resulting in a single gene that passed these selection steps. (D) 10-fold serial dilution spotting of cells expressing one of the toxins *mazF*, *relE*, or *hok* while co-expressing one of the random library hits (numbered 1-6) or an empty vector expressing the leader peptide only. Plasmids harbored the toxins under P*_ara_* or P*_van_* promoters as indicated. Plasmids carrying the random library hits driven by a P*_tet_* promoter. (E) Log_2_ fold-change of median msfGFP fluorescence levels of library hits 1-3 and hit 6 relative to the control strain with an empty vector expressing the leader peptide only. *msfGFP* expressed from either P*_ara_* or P*_van_*, as indicated. Data are the mean of two biological repeats.

We used this library to select for random genes that could enable cells to grow following induction of the toxin MazF, an endoribonuclease that degrades a wide range of cellular RNAs to inhibit cell growth^26^. We transformed our random gene library into an *E. coli* strain that expresses *mazF* from an arabinose-inducible promoter (P*_ara_*) on the chromosome. We then induced expression of both the random genes and *mazF* to select for those genes that inhibit MazF and promote growth. Because genomic mutations that prevent *mazF* expression, e.g. P*_ara_* mutations, also promote growth, plasmids in such backgrounds manifest as false positives. To enrich for true-positive hits, we performed two selection rounds; plasmids from the first round were harvested and used to transform new cells harboring P*_ara_*-*mazF* (Fig. 1C).

Deep sequencing of the library after two selection rounds against P*_ara_*-*mazF* revealed ∼2,000 enriched, random genes. We arbitrarily chose five of these random genes and tested whether each could inhibit two additional toxins: RelE, a ribonuclease^27^ of a different family than MazF, and Hok, a short hydrophobic toxin that depolarizes the cell membrane^28^. All five library hits could inhibit these toxins, which were also expressed from the arabinose-inducible promoter (Fig. 1D), and failed to inhibit MazF when the toxin was expressed from a vanillate-inducible promoter, P*_van_* (Fig. 1D). These observations suggest that these random genes are not directly inhibiting toxins, and instead somehow preventing transcription from the arabinose promoter. Consistent with this conclusion, we found that three of the random genes reduced the levels of monomeric, super-folding GFP (msfGFP) expressed from P*_ara_* and did not substantially affect msfGFP levels when expressed from the P*_van_* promoter (Fig. 1E).

To identify random genes that inhibit MazF independent of its expression system, we transformed the pool of ∼2,000 candidates into an *E. coli* strain in which *mazF* was expressed from P*_van_* (Fig. 1C). Two successive rounds of selection for growth on vanillate revealed a single random gene that could inhibit MazF driven by either P*_ara_* or P*_van_*, and that did not inhibit RelE or Hok (Fig. 1D). This gene did not affect levels of msfGFP produced from P_ara_ or P_van_ (Fig. 1E). We named this gene *ramF* for <u>r</u>andom <u>a</u>ntitoxin of <u>M</u>az<u>F</u>.

### RamF inhibits MazF by inducing its degradation

We sought to understand the molecular functions and adaptive advantages that the random protein RamF can provide cells. *ramF* has an open reading frame of 51 codons and an amino acid composition intermediate between small *E. coli* cytosolic and membrane proteins (Fig. 2A and Fig. S2). Searching in existing databases for proteins with sequence similarity to RamF did not return any significant matches. We first replaced the short ORF upstream of *ramF* with an RBS and confirmed the MazF-inhibitory activity of this new gene architecture (Fig. 2B). To confirm that the MazF-inhibitory activity of *ramF* depends on the production of a small protein, rather than mRNA, we mutated the start codon and found that this variant of *ramF* failed to inhibit MazF. We also generated a recoded variant of *ramF* with 46 synonymous mutations (representing changes to 30% of nucleotides in the ORF) and found that it could still inhibit MazF when co-expressed. Additionally, the originally selected *ramF* rescued growth inhibition following expression of a synonymously recoded *mazF* (83 mutations, 25% of the ORF) (Fig. 2B). Finally, *ramF* did not inhibit close homologs of the *E. coli* MG1655 *mazF*, the toxin used in our screen, as it did not rescue cells expressing *mazF* from the ECOR27 strain^29^ or MG1655 *chpB*, the closest *mazF* homolog in that strain (Fig. 2C). Taken together, these findings suggest that *ramF* encodes a novel protein that specifically alleviates the toxicity of the MG1655 MazF protein.

**Figure 2.**
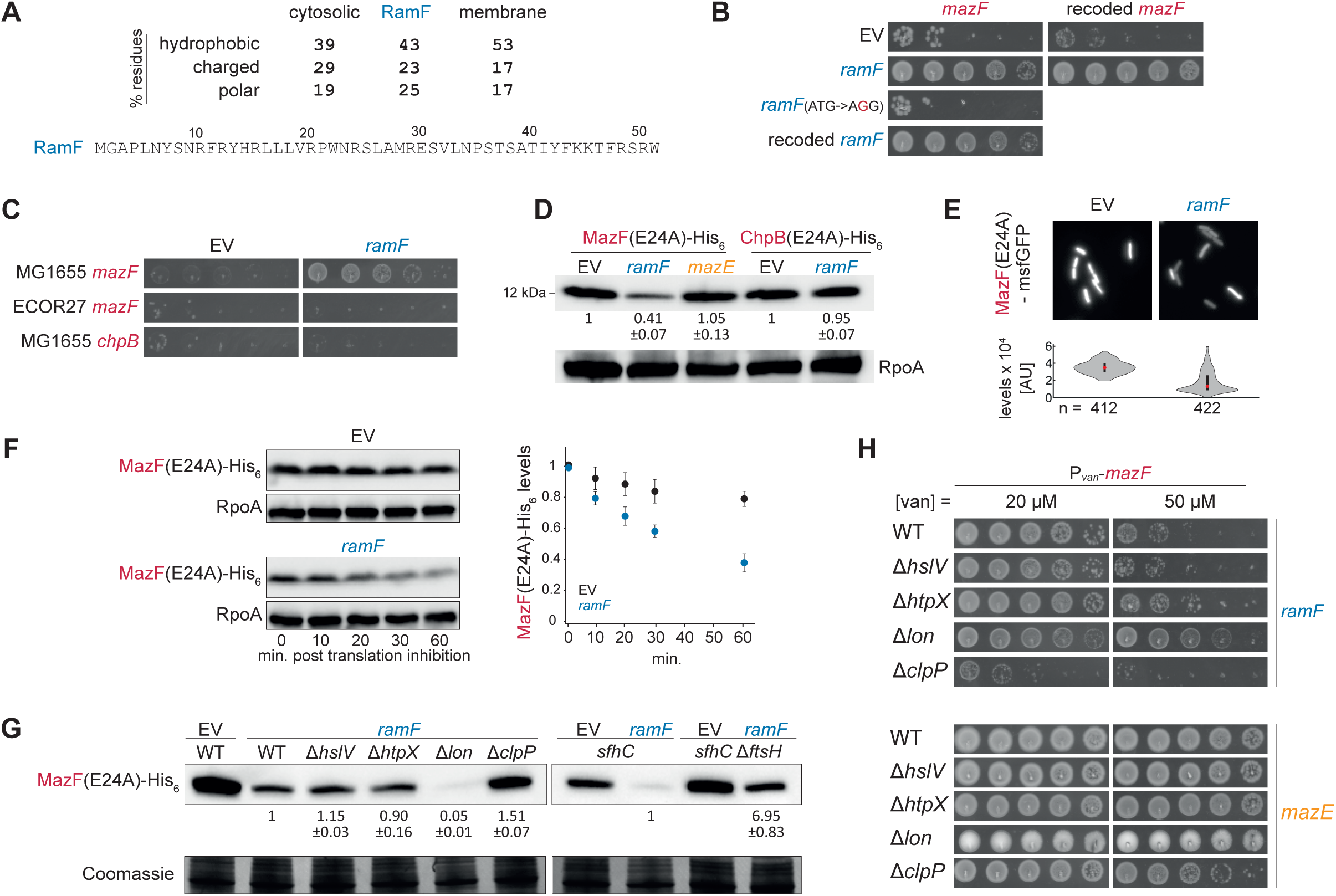
RamF is a specific-MazF inhibitor that induces MazF proteolysis. (A) Amino acid sequence of library hit 6, now named random <u>a</u>nti-toxin of <u>M</u>az<u>F</u> (*ramF*), and its amino acid composition compared to small proteins (< 100 amino acids) in *E. coli* that are either cytosolic (n=181) or membrane-localized (n=80). (B) 10-fold serial dilution spotting of cells expressing *mazF* with either its original nucleotide sequence or a synonymously recoded version from the P*_van_* promoter. Cells additionally expressed from the P*_tet_* promoter one of the following: *ramF*, *ramF* with a start codon mutation, synonymously recoded *ramF*, or an empty vector. The leader peptide of the library is not expressed in these cells or experiments hereafter. (C) 10-fold serial dilution spotting of cells expressing MG1655 *mazF* (reference sequence), ECOR27 *mazF* (56% identity), or MG1655 *chpB* (33% identity) from the P*_ara_* promoter. Cells also expressed *ramF* from the P*_tet_* promoter or carried an empty vector. (D) Immunoblot of MazF(E24A)-His_6_ or ChpB(E24A)-His_6_, expressed from P*_van_*, in cells co-expressing *ramF*, *mazE*, or harboring an empty vector. Quantification is based on three biological repeats and values are normalized to MazF(E24A)-His_6_ or ChpB(E24A)-His_6_ levels in the empty vector strains. (E) Fluorescence intensities of MazF(E24A)-GFP in cells expressing *ramF* or harboring an empty vector. At least 400 cells were measured for each strain. Violin plots: black bar represents the middle 50% of cells and red dot is the median. (F) Immunoblot of MazF(E24A)-His_6_, expressed from P*_van_*, from cells expressing *ramF* or harboring an empty vector. Time points were taken after the addition of tetracycline to stop the translation of new proteins. RpoA is a loading control. Quantification is based on three biological repeats and MazF(E24A)-His_6_ levels are normalized to t=0. (G) Same as (F) but for strains also lacking one of the major proteases of *E. coli*, as indicated. Loading control is based on Coomassie staining of total protein. Quantification for relevant strains is based on three biological repeats and values are normalized to MazF(E24A)-His_6_ levels in the *ramF*-expressing strain. Levels in the Δ*ftsH sfhC* strain are normalized to the *sfhC* control strain. (H) 10-fold serial dilution spotting of cells expressing *mazF* from P*_van_* and *ramF* or *mazE* from P*_tet_* in cells also lacking one of the major proteases of *E. coli*, as indicated. A plasmid harboring *mazF* could not be transformed into Δ*ftsH* cells.

How does RamF inhibit MazF? To test the effects of RamF on MazF levels, we generated a variant of MazF that could be easily used in molecular assays such as immunoblots as it harbored both a C-terminal His_6_-tag, which does not substantially impact function (Fig. S3A), and an E24A substitution, which was shown to reduce RNase activity of MazF but maintains a similar protein structure^30, 31^. Cells producing RamF had lower steady-state levels of MazF(E24A)-His_6_ compared to cells with an empty vector (Fig. 2D). Production of MazE, the natural antitoxin of MazF that inhibits its toxicity via direct binding^32^, did not reduce MazF levels (Fig. 2D). Producing RamF also reduced the fluorescence of MazF(E24A) fused to msfGFP in individual cells compared to a control strain (Fig. 2E, *P*<0.0001, t-test). These observations suggest that RamF inhibits MazF through a different mechanism than MazE, likely by reducing the toxin’s levels. RamF did not reduce levels of ChpB(E24A)-His_6_ (Fig. 2D), consistent with our finding that RamF did not neutralize ChpB toxicity (Fig. 2C).

Because RamF inhibits MazF in a promoter-independent manner, we hypothesized that RamF increases MazF degradation rather than reducing synthesis. To test this hypothesis, we treated cells producing MazF(E24A)-His_6_ with tetracycline to block new protein synthesis and followed MazF(E24A)-His_6_ levels by immunoblot over time. Cells producing RamF exhibited faster turnover of MazF(E24A)-His_6_ compared to control cells (Fig. 2F). This result indicates that RamF rescues MazF toxicity by promoting its degradation.

To identify the protease(s) that degrades MazF, we measured MazF(E24A)-His_6_ levels in strains producing RamF but lacking each of the major proteases in *E. coli* (Fig. 2G). Whereas the deletion of either *hslV* or *htpX* did not alter MazF(E24A)-His_6_ levels compared to wild-type cells, Δ*clpP* cells showed an increase in MazF(E24A)-His_6_ levels, suggesting that the ClpXP protease helps drive MazF degradation. Because *ftsH* is essential for viability, we could only examine the effects of Δ*ftsH* in the presence of the *sfhC* mutation^33^. Cells harboring Δ*ftsH* and the *sfhC* mutation showed significantly elevated levels of MazF(E24A)-His_6_ compared to an isogenic *sfhC* control, suggesting that FtsH plays a key role in MazF degradation. We also found that MazF(E24A)-His_6_ levels decreased in the Δ*lon* strain, possibly because ClpXP and FtsH are more active in the absence of Lon. Thus, we considered whether RamF inhibits Lon to drive increased degradation of MazF(E24A)-His_6_, thereby phenocopying the Δ*lon* strain. However, RamF did not decrease Lon activity (Fig. S5A-B), RamF could inhibit MazF in cells overproducing Lon (Fig. S5C), and producing the known Lon inhibitor PinA did no inhibit MazF (Fig. S5D). Thus, we concluded that RamF facilitates MazF-degradation through a Lon-independent mechanism.

Because the activity of RamF depends on toxin-induced degradation, we predicted that RamF inhibition efficiency would change in protease deletion strains that altered MazF levels. Indeed, for Δ*lon* cells in which MazF levels were reduced, RamF was functional at higher MazF induction levels than in control cells (Fig. 2H). In contrast, RamF did not inhibit MazF in Δ*clpP* cells as efficiently as in control cells (Fig. 2H), and it was impossible to transform a plasmid harboring *mazF* into Δ*ftsH* cells, presumably because even leaky expression leads to enough MazF accumulation that RamF cannot neutralize its toxicity. As controls, we confirmed that deleting either *hslV* or *htpX*, which did not affect MazF levels, did not affect RamF function. Additionally, we showed that the neutralization of MazF by MazE, which inhibits MazF independent of proteolysis, was not substantially affected by protease deletions.

### RamF interacts with chaperones to remodel protein homeostasis and cellular physiology

Our results demonstrated that RamF prevents MazF toxicity by facilitating its degradation, particularly via the FtsH protease. We found that known substrates of FtsH also exhibited decreased steady-state levels in RamF-producing cells (Fig. S6A), raising the hypothesis that RamF activates FtsH. However, overproducing FtsH in cells lacking RamF was insufficient to inhibit MazF and it did not alter RamF efficiency as a MazF inhibitor (Fig. S6B), suggesting that RamF does not inhibit MazF by simply activating FtsH.

How, then, can this randomly generated, 51-amino acid protein mediate MazF proteolysis? To characterize the physiological changes caused by RamF production, we first compared global RNA levels between cells expressing RamF to an empty vector control. We found that RamF does not lead to major transcriptional changes (Fig. 3A). There was, however, a ∼2.5-fold upregulation of the native *mazEF* locus (Fig. 3A left, red dots), which supports a model of RamF-dependent degradation of MazF because the complex MazEF negatively autoregulates *mazEF* expression^32, 34^; thus, degradation of MazF leads to up-regulation of *mazEF*. In agreement with RamF being a specific MazF inhibitor (Fig. 1 and 2), the mRNA levels of other TA systems, which are also autoregulated, were not affected (Fig. 3A left, pink dots, *P*=0.16, t-test).

**Figure 3.**
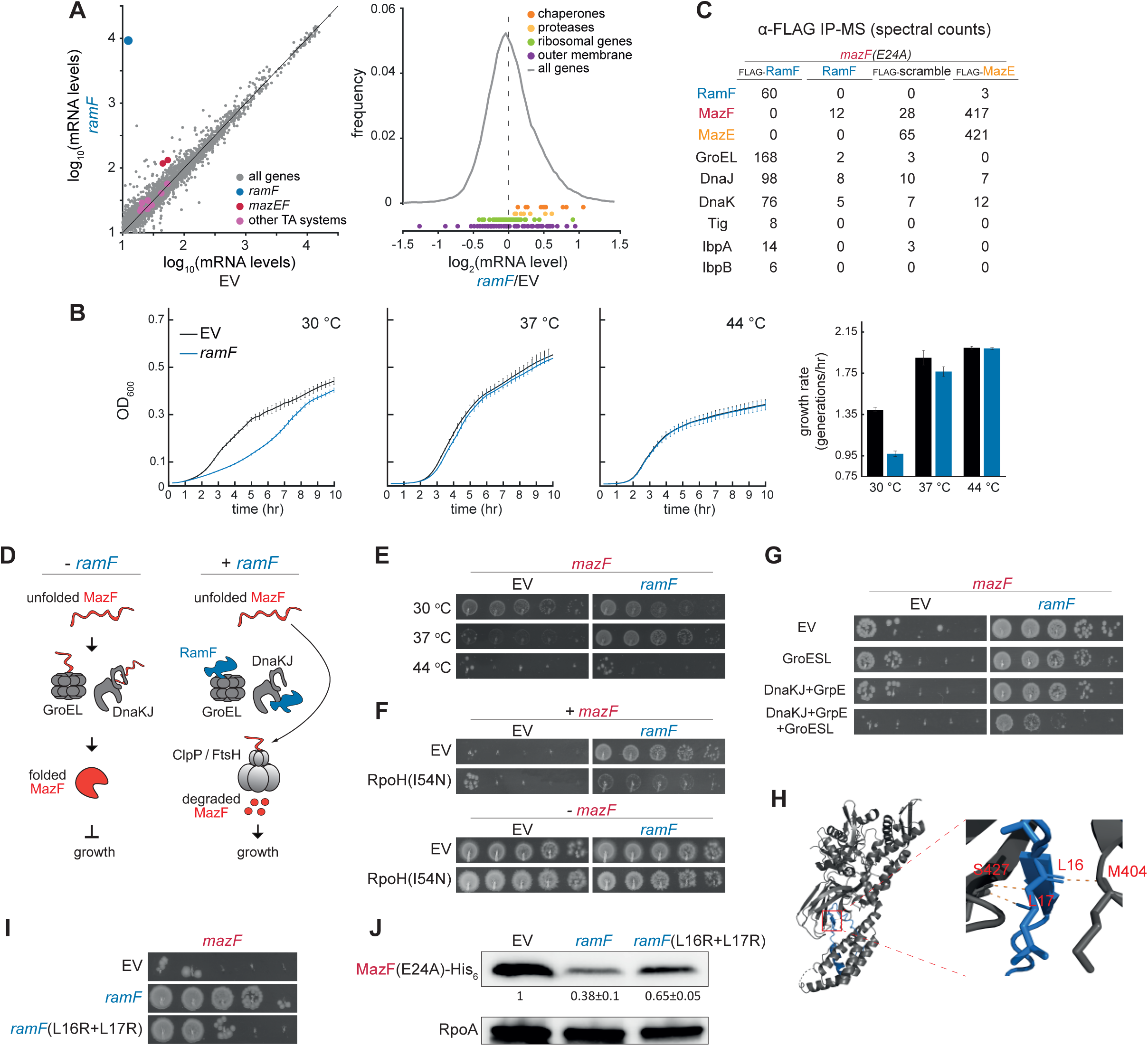
RamF interacts with chaperones to remodel protein homeostasis and cellular physiology. (A) (*Left*) Log_10_ of mRNA levels in Transcripts Per Million (TPM) for *E. coli* genes in cells expressing *ramF* or harboring an empty vector. (*Right*) Log_2_ of the mRNA level ratio between RamF-producing cells and cells with an empty vector. Colors: gray = all genes; red = *mazEF*; pink = other toxin-antitoxin systems; khaki = ribosomal protein genes; purple = outer-membrane genes; orange = chaperones; yellow = proteases. Data based on two biological repeats. (B) (*Left*) Growth curves for cells expressing *ramF* or harboring an empty vector growing at 30, 37, or 44 °C as a mean of at least 12 technical repeats. (*Right*) Maximal growth rate based on at least three biological repeats. (C) Spectral counts of *E. coli* proteins detected by mass-spectrometry following a pull-down with α-FLAG beads from a lysate of cells producing MazF(E24A) and (i) FLAG-RamF, (ii) RamF (negative control), (iii) FLAG-scrambled-RamF (negative control), or (iv) FLAG-MazE (positive control). (D) Model for RamF function as a MazF inhibitor. In cells not producing RamF, chaperones promote proper folding of MazF, leading to widespread RNA degradation and cell growth arrest. In RamF-producing cells, RamF binds chaperones and prevents MazF maturation, allowing FtsH and ClpXP to degrade MazF and restore cell growth. (E) 10-fold serial dilution spotting of cells expressing *mazF* from P*_ara_* and co-expressing *ramF* or harboring an empty vector, incubated at 30, 37, or 44 °C, as indicated. (F) 10-fold serial dilution spotting of cells expressing *mazF* from P*_van_*. Cells also express combinations of *ramF*, *rpoH*(I54N), or empty vectors, as indicated. (G) 10-fold serial dilution spotting of cells expressing *mazF* from P*_van_*. Cells also express combinations of *ramF*, *groESL*, *dnaKJ*+*grpE*, *groESL*+*dnaKJ*+*grpE*, or empty vectors, as indicated. (H) AlphaFold2 prediction of the interactions between residues M404 and S427 within the substrate-binding domain of DnaK with residues L16 and L17 of RamF. (I) 10-fold serial dilution spotting of cells expressing *mazF* from P*_van_* and co-expressing *ramF*, *ramF*(L16R+L17R), or an empty vector, as indicated. (J) Immunoblot of MazF(E24A)-His_6_, expressed from P*_van_*, from cells co-expressing *ramF*, *ramF*(L16R+L17R), or an empty vector. Loading control is RpoA. Quantification based on three biological repeats and values are normalized to MazF(E24A)-His_6_ levels in the empty vector strain.

Because RamF production results in MazF proteolysis, we tested if production of RamF affected protein homeostasis pathways, finding that chaperones and proteases were modestly, but significantly, up-regulated (Fig. 3A, right, *P*=1.94x10^-4^ and *P*=0.04, respectively, t-test). In comparison, the expression of other gene groups, *e.g.* ribosomal and outer-membrane gene groups, were unaffected by RamF (Fig. 3A, right, *P*=0.41 and *P*=0.21, respectively, t-test).

Our RNA-seq data suggest that RamF was well tolerated by cells and did not induce a strong stress response. In agreement, producing RamF had a minimal effect (0-2% reduction compared to control cells) on lag times and culture yields at 37 or 44 °C (Fig. S4). At 37 °C in LB medium, *ramF* expression only minimally reduced exponential-phase growth rate (Fig. 3B, *P*=0.04, t-test). At 44 °C, *ramF*-expressing cells grew identically to control cells (*P*=0.18, t-test), whereas at 30 °C *ramF* expression caused a severe growth reduction (*P*=1.27x10^-5^, t-test). This temperature-dependent phenotype further indicated that *ramF* activity may depend on protein homeostasis pathways as chaperone levels are often temperature-dependent^35–41^.

To further investigate how RamF affects cell physiology, we sought to find what proteins RamF interacts with in cells. To this end, we produced N-terminally FLAG-tagged RamF, which can still inhibit MazF (Fig. S3B) in cells also producing MazF(E24A). After immunoprecipitating RamF using α-FLAG beads, co-eluting proteins were identified by mass spectrometry. We did not detect MazF (Fig. 3C). As a control, we showed that the same procedure using a strain producing FLAG-MazE, did detect MazF, as expected. This result supports our conclusion that RamF inhibits MazF via a different mechanism than MazE.

Our mass spectrometry data revealed an enrichment of peptides from multiple chaperones, including GroEL (Hsp60), DnaK/J (Hsp70), trigger factor, and IbpA/B (Fig. 3C). In each case, the number of peptides found by immunoprecipitating FLAG-RamF was substantially higher than in two negative control experiments: a strain producing untagged RamF and a strain producing FLAG-tagged, but scrambled (same amino acid composition but in a randomized order) RamF protein that could not inhibit MazF (Fig. S3B).

In sum, our results demonstrated that RamF (i) drives increased proteolysis of MazF, (ii) promotes increased expression of chaperones and proteases, (iii) interacts *in vivo* with chaperones, and (iv) inhibits growth at a temperature where chaperone expression levels are relatively low. Based on these findings, we hypothesized the following molecular model for MazF inhibition by RamF: In cells lacking RamF, chaperones help MazF to adopt its native, folded state, which can then degrade RNA and thereby inhibit growth (Fig. 3D, left). In cells producing RamF, chaperones become occupied by RamF such that MazF is unable to fold properly, leaving it more susceptible to proteolysis, which allows cellular growth (Fig. 3D, right).

We next sought to test this model. First, we tested if temperature, which is correlated with chaperone levels, affects the ability of RamF to inhibit MazF. Indeed, we found that MazF failed to inhibit cellular growth at 30 °C even in the absence of RamF, possibly because of insufficient chaperone activity to fold MazF. Also, RamF rescued MazF toxicity at 37 °C, but not at 44 °C (Fig. 3E). These observations are consistent with our model, but growth temperature affects cell physiology in many ways. Thus, to modify chaperone availability in a more controlled manner, we used a strain producing a variant of the heat shock sigma factor (σ^32^) encoded by *rpoH* harboring an I54N substitution that prevents its degradation and thus maintains its activity^42^. RamF failed to rescue cells producing both MazF and RpoH(I54N), and this phenotype was not caused by toxicity of RpoH(I54N) overproduction, as cells producing it in the absence of MazF grew similarly to control cells (Fig. 3F). We also generated cells that overproduce either the chaperone system DnaK/DnaJ/GrpE or GroEL/GroES, or both. Overproducing the individual chaperone systems partially reduced the ability of RamF to alleviate MazF toxicity, with a substantial drop in RamF activity when overproducing both systems (Fig. 3G). Taken together, our results demonstrate that cellular availability of chaperones is critical to RamF function.

Finally, we asked if the interaction between RamF and chaperones detected in our IP-MS data is important for MazF inhibition. Using AlphaFold2^43, 44^, we modeled the interaction between RamF and DnaK and found that positions M404 and S427 in DnaK are predicted to bind positions L16 and L17 in RamF, respectively (Fig. 3H). Notably, these predicted DnaK positions were (i) found in the substrate-binding domain of DnaK^45^ and (ii) previously shown in a crystal structure to bind two contiguous Leu residues of a model peptide^46, 47^. A variant of RamF with the substitutions L16R and L17R inhibited MazF less efficiently (Fig. 3I) and induced MazF degradation less efficiently than RamF, consistent with our model that RamF’s interaction with chaperones is critical to MazF inhibition.

### RamF specificity towards MazF is partially determined by the N terminus of the toxin

Our results thus far indicate that RamF interacts with central protein homeostasis pathways, which ultimately results in MazF proteolysis. Given this function, how does RamF inhibit *E. coli* MG1655 MazF but not other close MazF homologs (Fig. 1B)? We speculated that this specificity might stem from *E. coli* MG1655 MazF, but not its homologs, being recognized by FtsH. The FtsH protease has been shown to recognize substrates by unique degron sequences found at the N or C terminus of proteins, or internally^48–51^. Because C-terminal tagging of MazF did not change RamF-dependent inhibition (Fig. S3B), we tested the relevance of its N terminus to degradation. We fused an N-terminal myc tag to MazF and found that while inhibition by MazE was maintained, the tag abolished inhibition by RamF (Fig. 4A). This result suggests that tagging MazF on its N terminus prevented its degradation, presumably by occluding the degron. Indeed, myc-MazF(E24A)-His_6_ levels did not drop in cells expressing *ramF*, in contrast to the levels of MazF(E24A)-His_6_ (Fig. 4B). Additionally, we found that removing amino acids 2-6 or 2-10 eliminated MazF toxicity (Fig. 4C), suggesting that this region not only mediates MazF degradation but is also essential to the ability of MazF to mature and perform its toxic function.

**Figure 4.**
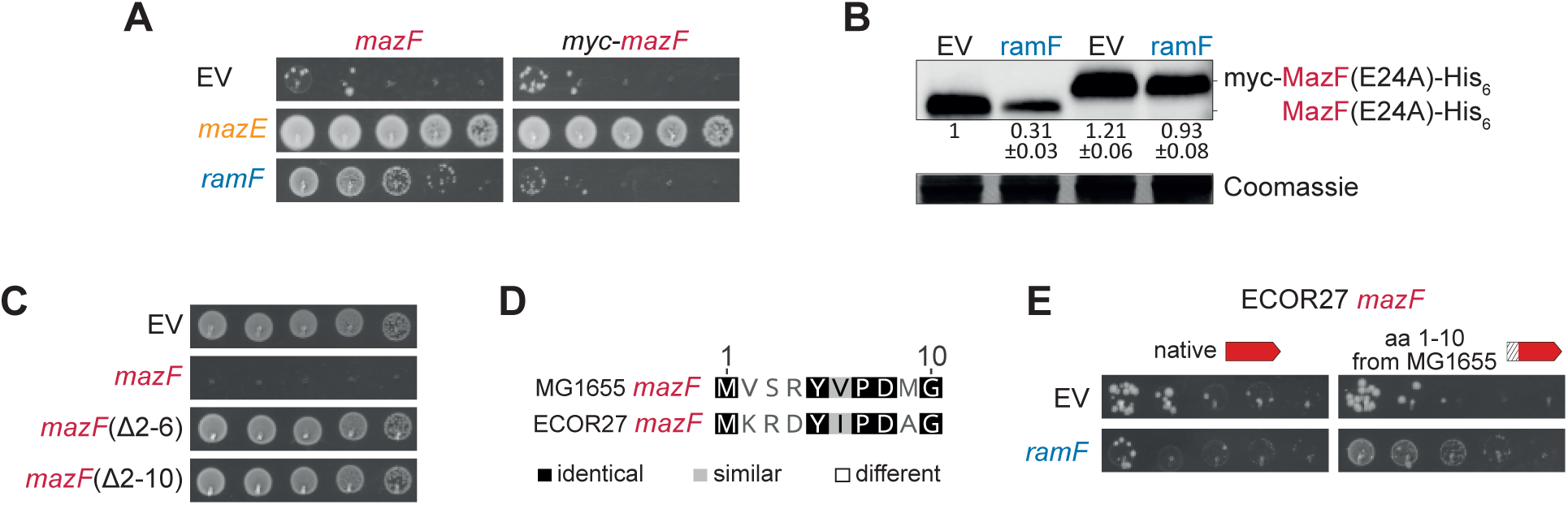
The N terminus of MazF is essential for its inhibition by RamF. (A) 10-fold serial dilution spotting of cells expressing *mazF* or *myc-mazF* from P*_van_*. Cells also express *ramF*, *mazE*, or an empty vector, as indicated. (B) Immunoblot of MazF(E24A)-His_6_ or myc-MazF(E24A)-His_6_, expressed from P*_van_*, from cells co-expressing *ramF* or harboring an empty vector. Loading control is based on Coomassie staining of total protein. Quantification based on three biological repeats and values are normalized to MazF(E24A)-His_6_ levels in the empty vector strain. (C) 10-fold serial dilution spotting of cells expressing *mazF*, *mazF*(Δ2-6), *mazF*(Δ2-10), or an empty vector, as indicated. (D) Sequence alignment of the first ten positions of MG1655 MazF and ECOR27 MazF. (E) 10-fold serial dilution spotting of cells expressing ECOR27 *mazF* or ECOR27 *mazF*(1-10 from MG1655) from P*_ara_*. Cells are additionally expressing *ramF* or an empty vector, as indicated.

A sequence alignment of MG1655 MazF and ECOR27 MazF indicated that the first ten amino acids differ at five positions (Fig. 4D, see full alignment in Fig. S7). We hypothesized that replacing these amino acids in ECOR27 MazF with those of MG1655 MazF might make this chimeric protein a better FtsH substrate and therefore sensitive to RamF inhibition. Indeed, RamF gained the ability to inhibit ECOR27 MazF when its first ten amino acids matched those in MG1655 MazF (Fig. 4E). Taken together, our results explain how a novel, random protein that interacts with central cellular pathways can (i) have a specific effect on a single target and (ii) how accumulation of mutations on new targets can make them susceptible to this effect.

### Beneficial mutations that optimize the function of RamF as a MazF inhibitor are common

Once a *de novo* gene like *ramF* is established in a genome, natural selection can, in principle, further improve its activity via subsequent beneficial mutations. To ask whether RamF can become a better MazF inhibitor, we used PCR-based mutagenesis to create a library of ∼60,000 RamF variants. This library was then transformed into the same *E. coli* strain used in the initial screen and selected on higher levels of MazF such that MazE rescues growth, but the original RamF cannot (Fig. 5A-B, see Methods). The library was deep-sequenced pre- and post-selection to find mutations that were enriched following selection (Fig. 5C).

**Figure 5.**
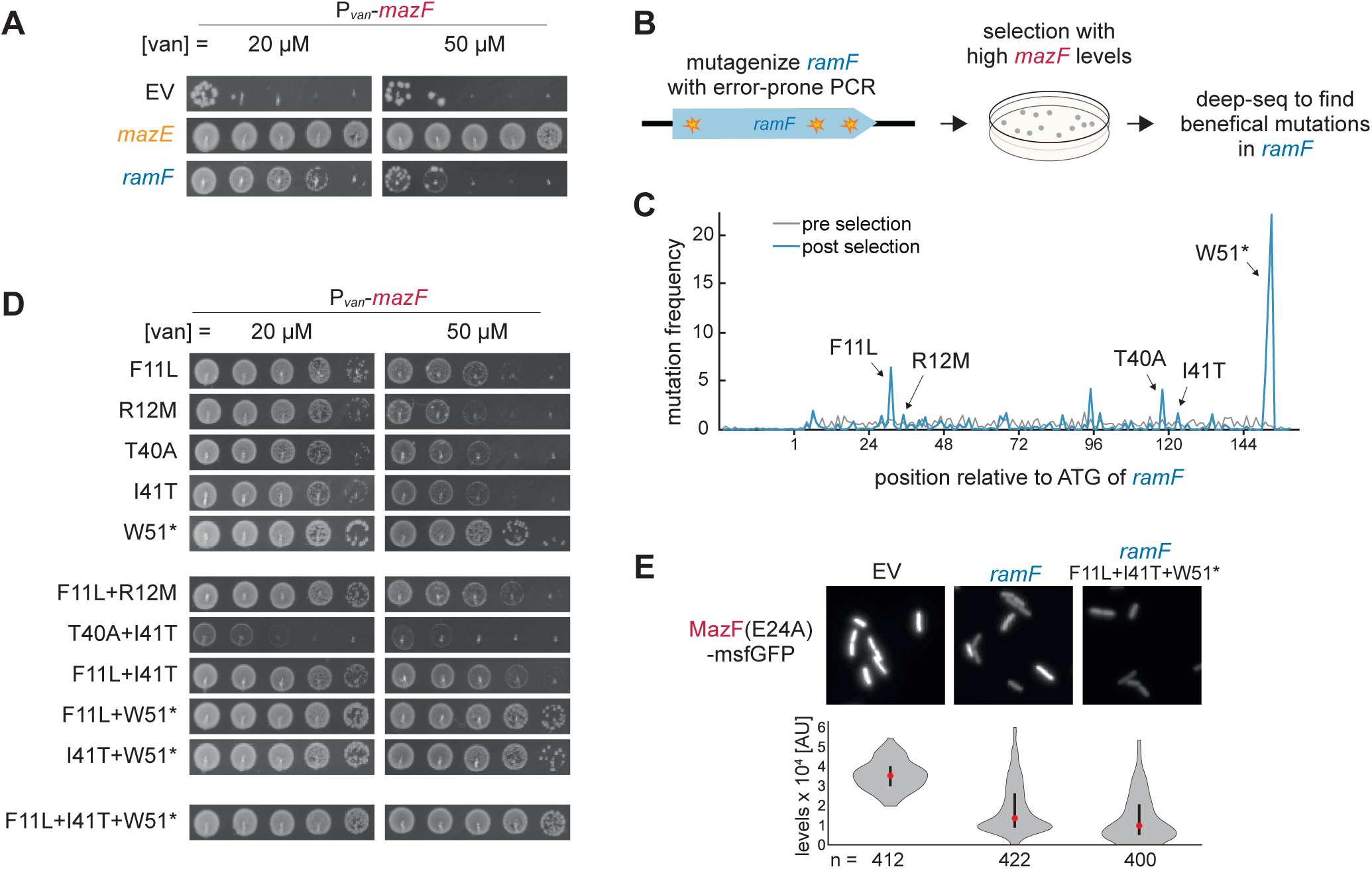
Beneficial mutations that optimize the function of RamF as a MazF inhibitor. (A) 10-fold serial dilution spotting of cells expressing *mazF* from P*_van_*. Cells are additionally expressing *ramF* or *mazE*, as indicated. (B) Selection strategy for identifying beneficial mutations that improve RamF activity. Variants of *ramF*, generated by random mutagenesis of *ramF* with error-prone PCR, were selected in the presence of high MazF levels that the original RamF cannot neutralize. (C) Frequency of *ramF* variants pre- and post-selection on high MazF levels. (D) 10-fold serial dilution spotting of cells expressing *mazF* from P*_van_*. Cells were additionally expressing *ramF* variants, as indicated. (E) Data as in Fig. 2E, with the addition of RamF(F11L+I41T+W51*)-producing cells.

We found five mutations that individually improved RamF’s inhibition of MazF spread across three regions along the *ramF* ORF: F11L, R12M, T40A, I41T, and W51* (Fig. 5D). Combinations of these mutations mostly showed additive phenotypes, except for the substitutions T40A and I41T which exhibited strong negative epistasis (Fig. 5D). We also generated an improved RamF variant harboring F11L, I41T, and W51*, which was the most efficient MazF inhibitor (Fig. 5D). We confirmed that the RamF(F11L+I41T+ΔW51) variant also reduced MazF(E24A)-msfGFP levels further compared to cells expressing RamF (Fig. 5E, *P*<0.0001, t-test). Taken together, our results suggests that beneficial mutations that improve RamF functions are common and easily accessible by natural selection.

## Discussion

There has been a growing interest recently in studying the evolutionary origin, phylogenetic distribution, and functions of small proteins (< 50 aa) in biological systems^52–54^. New detection methods, based on genome annotation algorithms or experimental techniques like mass spectrometry and ribosome profiling, have improved the identification of small ORFs^55–59^, which are abundant across the tree of life. While the functionality of most small proteins is still an open question, several have been shown in bacteria to interact with and regulate protein kinases, transporters, metabolic enzymes, and cell division proteins in the cytosol and the membrane^60^.

The evolutionary origin of small proteins is also unclear. Did the small genes that encode them evolve from duplicated fragments of longer genes, or did they emerge *de novo* from promiscuous, yet beneficial expression of random ORFs? This is a tricky question to answer because of the small size of these ORFs, which poses a challenge for homology detection. However, synteny-based approaches have identified some genes that almost surely arose *de novo* in previously intergenic regions^6, 61, 62^. Additionally, there is ample evidence for the spurious expression of small RNAs and proteins^7, 11–15^, many of which may have neutral fitness consequences, providing evolution with a pool of candidates for novel genes^13^.

A complementary approach to studying *de novo* genes is the screening of synthetic, random proteins for functions. Such experiments could support a *de novo* origin of small proteins by showing how random proteins, not previously present in cells, have the potential to integrate beneficially within cells. *In vitro* screens have identified some functions for random proteins, including the ability to bind ATP^21^, lipopolysaccharides^22^, or streptavidin^19^. However, these binding abilities have not been demonstrated *in vivo*, and the toxicity of these peptides to cells has not been tested. Another study showed how different chaperones can change the solubility of random proteins in a cell-free expression system^17^, but did not explore the *in vivo* effects of chaperones on random proteins.

In a few cases, random protein libraries have been screened *in vivo*. One recent study reported that random proteins in *E. coli* can have beneficial effects on growth^63^, but serious caveats in experimental design were subsequently raised^64, 65^. An additional study found random proteins that rescue growth of an *E. coli* serine auxotroph^66^. However, these random proteins strongly activated the cellular SOS response, suggesting that their production has a fitness cost. Another random protein was found to promote survival in high concentrations of copper^67^, but the underlying molecular mechanism remains unknown. This random protein was also studied *in vitro* and found to catalyze the conversion of cysteine to H_2_S in a pyridoxal phosphate dependent manner^20^.

Two additional studies found hydrophobic proteins that insert into the cell membrane to provide antibiotic resistance to *E. coli* cells^23, 24^. In one case, the small protein depolarizes the membrane, which protects against antibiotics that require a strong proton motive force for cell entry, but this small protein severely diminished cell growth both in the presence and absence of antibiotics. In the second case, a random protein that stimulates a membrane-bound histidine kinase was identified, but it substantially increased growth lag time and decreased culture yield by 10-20%. In sum, there has been limited evidence of random proteins that can be stably produced *in vivo* and provide a benefit to cells through a cytosolic function without substantially affecting cell growth.

Here, we identified and characterized a random protein, RamF, that does not substantially impact growth lag time, growth rate, or culture yield, particularly at 37 °C, the temperature at which RamF was selected. RamF interacts with cytosolic chaperones, which remodels the physiology of *E. coli* cells to promote their survival in the presence of MazF. Because we know that *ramF* originated from a *bona fide* random sequence, our findings support the idea that natural, small proteins that function in the cytosol may have originated from expression of random sequences to modulate pre-existing cellular pathways in response to an evolutionary challenge.

RamF was our only hit from a pool of ∼10^8^ sequences that inhibited MazF by affecting the toxin’s protein levels rather than *mazF* transcription. Given the tendency of random proteins to include hydrophobic regions^5, 16^ and bind chaperones *in vitro*^17^, why did we not observe more hits that work similarly? While there are probably many other hydrophobic random proteins in our library, they likely suffer from one or more of the following shortcomings: (i) they are not stable, with a fast turnover that prevents functional interactions with cellular components, (ii) they have a transmembrane domain and are localized to the membrane, (iii) their hydrophobic amino acid composition leads to toxic aggregation, or (iv) they activate the stress responses that offset any beneficial change in cell physiology that these potential hits may have. Only random proteins that avoid these hurdles might modify cells in a beneficial way that promotes MazF proteolysis.

For a random gene to emerge and be retained, it must provide a beneficial function to cells. Once established, such a gene can improve via beneficial mutations. We found five beneficial mutations that make RamF a better MazF inhibitor, suggesting that random proteins can be better integrated into cellular systems as they evolve. How these mutations individually, or in combination, improved RamF function is an open question for future exploration.

We screened a library of random proteins against the toxin MazF, but when do biological systems face this challenge? Toxin-antitoxin (TA) systems are widespread in bacteria and found on both chromosomes and plasmids^68, 69^. Notably, the antitoxins for the homologs of a given toxin are often not homologous suggesting that antitoxins can readily change, and possibly arise *de novo* via a pathway similar to that reported here for RamF. Additionally, antitoxins are often short proteins harboring unstructured domains, which bind their toxin counterparts^70–74^. These qualities - short length and containing unstructured domains - are also common among random sequences and young genes^16, 75, 76^, further supporting the possibility that antitoxins have arisen *de novo*.

Short proteins with a similar antitoxin function to RamF might be found in phages. A growing body of work suggests that TA systems, including *mazEF* system^77^, provide anti-phage defense to bacterial cells via an abortive infection mechanism^78–81^. Upon cellular entry of phage, toxins are liberated from their antitoxins, often leading to cell death, which prevents phage replication and spread through a population of cells. Interestingly, phages can ‘fight back’, sometimes by producing small proteins that inhibit toxins and thus allow phages to complete their infection cycle^82, 83^. Homologs of these short proteins are often only found in phage genomes^84^, and our work suggests a possible *de novo* origin for these proteins.

Finally, we identified ∼2,000 hits that block MazF in a promoter-dependent manner likely by reducing expression from the P*_ara_* promoter, though we have not characterized the molecular functions of these genes in full. Why did we find significantly more hits that presumably target the activity of the arabinose promoter than inhibit MazF itself? A likely explanation is that the complex arabinose pathway in *E. coli*, including the arabinose uptake machinery, metabolic enzymes, and regulatory proteins^85–87^ provides more opportunities for random proteins to directly or indirectly prevent activation of the arabinose-inducible promoter. Additionally, inhibiting the arabinose pathway may be less likely to perturb essential cellular functions, allowing more solutions using this mechanism to emerge. Whatever the case, the P*_ara_*-dependent hits serve as additional evidence that random proteins can readily adopt beneficial functions inside cells. The idea that expression of random sequences, probably through spurious transcription and translation in natural, biological systems, can be advantageous is critical for *de novo* genes to emerge. Our work demonstrates the feasibility of this randomness-to-function process and provides molecular insight into how *de novo* genes can integrate into existing cellular pathways.

## Materials & Methods

### Plasmids, strains, and growth conditions

All strains and plasmids used in this study are listed in Table S1. *Escherichia coli* was grown in LB medium (10 g/L NaCl, 10 g/L tryptone, 5 g/L yeast extract) or M9 medium (10x stock made with 64 g/L Na_2_HPO_4_-7H_2_O, 15 g/L KH_2_PO_4_, 2.5 g/L NaCl, 5.0 g/L NH_4_Cl supplemented with 0.1% casamino acids, 0.4% glycerol, 2 mM MgSO_4_, and 0.1 mM CaCl_2_). In cases M9 was used, 0.8% glucose was added to prevent leaky expression from the arabinose-inducible promoter. Media for selection or plasmid maintenance were supplemented with carbenicillin (100 μg/mL), chloramphenicol (20 μg/mL), or kanamycin (30 μg/mL) as appropriate. Overnight cultures were prepared in the same medium used in a given experiment and cells were grown at 37 °C and 180 rpm in an orbital shaker. The arabinose-, tetracycline-, and vanillate-inducible promoters were induced with 0.0002%-0.2% arabinose, 100 ng/μL anhydrous tetracycline (aTc), 15-100 μM vanillate, respectively.

Plasmids were generated by Gibson assembly according to the manufacturer’s protocol. Inserts were either amplified from a template by PCR or commercially synthesized by Integrated DNA Technology as gBlocks. All plasmids were confirmed by Sanger sequencing of the inserts or by full-length plasmid sequencing by Plasmidsaurus. Plasmids were introduced into cells by either TSS transformation or electroporation.

### *E. coli* genome engineering

To construct *E. coli* BW27783 *amyA*::P_ara_-*toxin/msfGFP* (strains ML-4045 to ML-4048) and *E. coli* BW27783 *amyA*::P_van_-*toxin/msfGFP* (strains ML-4049 to ML-4050), the “P_ara_-*toxin*, *kan^R^*” or “P_van_-*toxin*, *kan^R^*” cassettes were PCR amplified from plasmids with primers that included homology to the *amyA* locus. These amplicons were inserted into the genome of the arabinose titratable strain BW27783^88^ using the lambda Red-based recombination^89^. Single insertions were confirmed by PCR and Sanger sequencing for individual colonies.

### Assembly and transformation of the random gene library

The random gene library was constructed by cloning 150 random nucleotides into the vector ML-4052 such that they immediately followed an ATG and were followed by two TAA stop codons. Specifically, pooled ssDNA oligos of 50 NNB codons flanked on their 5’ end by the sequence GCCTGGCTACCGTCTCGTATG and on their 3’ end by TAATGGAGACGAGCAGGCGATG were synthesized by Integrated DNA Technologies (IDT). To avoid frequent premature stop codons, NNB codons, rather than NNN codons, were used; NNB libraries produce similar amino acid composition as NNN libraries. Oligos were PCR amplified using KAPA enzyme according to manufacturer recommendations with 16 amplification cycles. Six independent reactions were performed and combined to minimize PCR bias. Amplicons of the expected size of 193 nucleotides were purified from a gel using a Zymo Gel DNA Recovery kit and ∼500 ng of this insert dsDNA were digested and cut using the type IIS restriction enzyme Esp3I at 37 °C for 3 hours to reach full digestion. Approximately 500 ng of the vector ML-4052 were similarly cut by BsmBI, and both the insert and vector were subsequently purified on a Zymo DNA clean column. Then, 250 femtomol of the vector and 1.25 pmol of the insert were combined in a 20 µL ligation reaction with T4 ligase and Esp3I enzyme. The ligation reaction was cycled between 16 °C for 2 min and 37 °C for 2 min for 100 cycles to allow iterative ligation and digestion. This approach increased the ligation efficiency because once an insert was ligated to a vector it could no longer be cut by the restriction enzyme. Ligations were dialyzed on Millipore VSWP 0.025 μm membrane filters for 60 min and then the entire volume was electroporated into 20 μL of Invitrogen MegaX DH10B cells, which resulted in ∼10^8^ transformants. Transformants were grown overnight (14 h) in 50 mL of LB + carbenicillin. Then, the culture was split: 25 mL were frozen in 20% glycerol for long-term storage at -80 °C and 25 mL were prepped for plasmids. The plasmid library of random genes was then dialyzed and electroporated into *E. coli* strain ML-4045 to yield ∼5 × 10^8^ transformants.

### Amplicon sequencing of random library and analysis

To assess the library complexity pre- and post-selection, random sequences were amplified using a forward primer that included the Illumina anchors and indexes as well as a region directly upstream of the random nucleotides and a reverse primer matching a region immediately downstream of the random nucleotides. PCR reactions were performed using KAPA enzyme according to manufacturer recommendations with 10 amplification cycles. Four independent reactions were performed and combined to minimize PCR bias. Amplicons were purified from an agarose gel using a Zymo Gel DNA Recovery kit. Paired-end sequencing was performed on an Illumina MiSeq at the MIT BioMicroCenter. Paired-end reads were merged using PEAR with default parameters, and identical reads were clustered using usearch with default parameters.

### Bacterial growth by spotting assay on solid media

In experiments with P_ara_ induction, cultures were grown to saturation overnight in M9-glucose supplemented with 5% LB and the appropriate antibiotics. Cultures were then serially diluted tenfold and spotted on appropriate plates supplemented with 0.8% glucose (toxin repressing), 0.0002%-0.2% arabinose (toxin inducing), 100 ng/μL aTc (random gene inducing), or 0.0002%-0.2% arabinose and 100 ng/μL aTc (toxin and random gene inducing). Plates were then incubated at 37 °C for 24-36 hr before imaging. A similar approach was used in experiments with P_van_ induction, only that LB medium and 15-100 μM vanillate as inducer were used.

### Bacterial growth in liquid

Cultures were grown overnight at 30 °C in an appropriate medium, back-diluted 1:50, and grown an additional overnight at 30 °C. The next day cultures were diluted 1:200 and seeded into a 96-well plate (160 µL culture overlaid with 70 µL mineral oil) such that each culture had 12 replicates on the same plate and plates were replicated independently at least three times. Growth was monitored at 15 min intervals with orbital shaking on a plate reader (Biotek) at the indicated temperature. Data presented are the mean and standard deviation of all replicates.

### Measurements of msfGFP levels with flow cytometry

Strain ML-4048 or ML-4050 with plasmids ML-4052 to ML-4055 or ML-4058 were grown overnight at 37 °C in LB supplemented with appropriate antibiotics. Cultures were diluted 1:500 in medium supplemented with 100 ng/μL aTc to induce expression of the random genes (or an empty vector) and grown for 30 minutes at 37 °C. Then, either 0.2% arabinose or 100 μM vanillate was added to induce the expression of msfGFP. Cultures were grown an additional 4.5 hours at 37 °C, then diluted 1:40 into PBS supplemented with a high concentration of (0.5 g/L) kanamycin to stop translation, and incubated at room temperature for 10 min. Fluorescence was measured on a Miltenyi MACSQuant VYB. Two independent cytometry experiments were performed for each strain and 30,000 cells were measured per replicate. FlowJo was used to analyze the data, gating on single live cells, and extracting the median of the msfGFP distribution.

### Western blot analysis of steady-state MazF(E24A)-His_6_ levels

Cultures were grown overnight at 37 °C in an appropriate medium, back-diluted 1:200 the next day, and grown at 37 °C until OD_600_ ∼ 0.2. Then, 100 ng/μL aTc was added to induce *ramF* (or an empty vector) and cultures were grown for an additional 30 min. Next, 100 µM vanillate was added to induce *mazF(E24A)-His_6_*, and cultures were grown for an additional 60 min. At that point, 1 mL of cells was pelleted and flash-frozen. Pellets were then resuspended in 1× Laemmli sample buffer (Bio-Rad) supplemented with β-mercaptoethanol normalized to the OD_600_ of the culture at the moment of collection. Samples were boiled at 95 °C for 5 min, analyzed by 4%-20% SDS–PAGE, and transferred to a 0.2 μm PVDF membrane. To visualize proteins, an anti-His_6_ antibody (ThermoFisher) was used at a final concentration of 1:1000, and SuperSignal West Femto Maximum Sensitivity Substrate (Invitrogen) was used to develop the blots. Blots were imaged by a ChemiDoc Imaging system (Bio-Rad). Images shown are one of at least three independent biological replicates. Band intensities were quantified using ImageJ (https://imagej.nih.gov/ij), and averages and standard errors are based on all replicates. Loading controls were performed using either an anti-RpoA antibody (Biolegend) at a final concentration of 1:5000 or a Coomassie stain as previously described^90^.

### MazF degradation assay

Cultures were grown overnight at 37 °C in an appropriate medium, back-diluted 1:200 the next day, and grown at 37 °C until OD_600_ ∼ 0.2. Then, 100 µM vanillate was added to induce *mazF(E24A)-His_6_* and cultures were grown for an additional 60 min. Next, 100 ng/μL aTc was added to induce *ramF* (or an empty vector) and cultures were grown for an additional 30 min. At that point, 1 mL of cells was pelleted and flash-frozen. 100 µg/mL Tetracycline was added to block protein synthesis and samples were collected at time points 10, 20, 30, and 60 min. Immunoblots for samples were performed as described above, using RpoA as a loading control.

### Immunoprecipitation-Mass Spectrometry (IP-MS)

*E. coli* strains with plasmids ML-4060, ML-4075, ML-4076, or ML-4078 were grown overnight in LB supplemented with appropriate antibiotics at 37 °C. Overnight cultures were back-diluted 1:200 in 50 mL and grown until OD_600_ ∼ 0.2 at 37 °C. Then, 100 ng/μL aTc was added to induce FLAG-RamF or RamF or FLAG-scrambled RamF or FLAG-MazE, and cultures were grown for an additional 30 min. Next, 100 µM vanillate was added to induce MazF(E24A) and cultures were grown for additional 60 min. Cultures were pelleted at 4,000 *g* for 10 min at 4 °C, supernatant was removed, and cells were resuspended in 900 μL lysis buffer (B-PER II, ThermoFisher) supplemented with protease inhibitor (Roche), 1 μL/mL Ready-Lyse Lysozyme Solution (Lucigen) and 1 μL/mL benzonase nuclease (Sigma). Samples were incubated at room temperature for 15 min, normalized by OD_600_, and centrifuged at 15,000 × g for 20 mins at 4 °C. Next, 850 μL of supernatant were incubated with pre-washed anti-FLAG M2 magnetic beads (Sigma) for 1 h at 4 °C with end-over-end rotation after which beads were washed 3 times with a wash buffer free of detergent (25 mM Tris-HCl, 150 mM NaCl, 1 mM EDTA, and 5% glycerol). On-bead reduction, alkylation, and digestion were performed. Proteins were reduced with 10 mM dithiothreitol (Sigma) for 1 h at 56 °C and then alkylated with 20 mM iodoacetamide (Sigma) for 1 h at 25 °C in the dark. Proteins were then digested with modified trypsin (Promega) at an enzyme/substrate ratio of 1:50 in 100 mM ammonium bicarbonate, pH 8 at 25 °C overnight. Trypsin activity was halted by the addition of formic acid (99.9 %, Sigma) to a final concentration of 5%. Peptides were desalted using Pierce Peptide Desalting Spin Columns (Thermo) and then lyophilized. The tryptic peptides were subjected to LC–MS/MS. Peptides were separated by reverse phase HPLC (Thermo Ultimate 3000) using a Thermo PepMap RSLC C18 column over a 90-min gradient before nano-electrospray using an Exploris mass spectrometer (Thermo). Solvent A was 0.1% formic acid in water and solvent B was 0.1% formic acid in acetonitrile. Detected peptides were mapped to *E. coli* MG1655 protein sequences with the addition of the RamF sequence, and protein abundance was estimated by the number of spectrum counts. For full IP-MS results of each pull-down, see Table S2.

### RNA extraction and sequencing

*E. coli* strains with plasmids ML-4059 or ML-4060 were grown overnight in LB supplemented with appropriate antibiotics at 37 °C. Overnight cultures were back-diluted 1:200 in 25 mL cultures and grown until OD_600_ ∼ 0.2 at 37 °C. Then, 100 ng/μL aTc was added to induce RamF or empty vector and cultures were grown for an additional 45 min. At that time, 1 mL of each culture was mixed with stop solution (110 µL; 95% ethanol and 5% phenol) and pelleted by centrifugation for 30 sec at 16,000 x g on a tabletop centrifuge. Pellets were flash-frozen and stored at -80 °C. Cells were lysed by adding TRIzol (Invitrogen) preheated to 65 °C directly to pellets, followed by 10 min of shaking at 65 °C and 2,000 rpm on a ThermoMixer (Eppendorf). RNA was extracted from the TRIzol mixture using Direct-zol (Zymo) columns according to manufacturer’s protocol. Genomic DNA was removed by adding 2 µL of Turbo DNase (Invitrogen) in a 100 µL final volume using the provided buffer and incubating for 30 minutes at 37 °C. DNase reaction products were cleaned up with a Zymo RNA clean and concentrator kit and eluted in 25 µL water.

Libraries were generated as described previously^26^. The library generation protocol was a modified version of the paired-end strand-specific dUTP method using random hexamer primers. rRNA was removed using a recently-developed do-it-yourself *E. coli* rRNA depletion kit, using 2.5 mg total RNA as input^91^. Paired-end sequencing was performed on an Illumina MiSeq at the MIT Bio Micro Center.

Geneious Prime 2022.2.2 used to map reads to the *E. coli* MG1655 genome (accession number NC_000913) with default parameters and to calculate Transcripts Per Million (TPM) values for all genes. TMP values of each sample were normalized by the median TMP value of a given sample to make all samples comparable^92^. Data shown are based on two independent repeats for each strain. Raw data can be found with NCBI BioSample accessions SAMN32730695 and SAMN32730696.

### Microscopy

*E. coli* strains with plasmid ML-4093 and additional plasmids ML-4059, ML-4060, or ML-4074 were grown in LB supplemented with appropriate antibiotics overnight at 37 °C. Cultures were diluted 1:200, grown at 37 °C for 30 min, supplemented with 100 ng/μL aTc to induce RamF or empty vector, and cells were grown for additional 30 min. Next, 0.2% arabinose was added to induce msfGFP, and cells were grown for 2.5 hours at 37 °C. 1 μL of each culture was spotted onto a 1% agarose pad prepared with PBS and placed in a 35 mm glass-bottom dish with 20 mm microwell #0 coverglass (Cellvis). Phase-contrast and epifluorescence images were taken using a Hamamatsu Orca Flash 4.0 camera on a Zeiss Observer Z1 microscope using a ×100/1.4 oil immersion objective and an LED-based Colibri illumination system using MetaMorph software (Molecular Devices). Images were analyzed in Fiji using the MicrobeJ plug-in^93^. Individual cells were identified by the phase-contrast image and fluorescence intensity was recorded for each cell, with at least 400 cells for each culture.

### Error-prone PCR mutagenesis of RamF

RamF was mutagenized using error-prone PCR-based mutagenesis, as previously described^94^. *ramF* was amplified using Taq polymerase (NEB) and 0.5 mM MnCl_2_ was added to the reaction as the mutagenic agent. PCR products were treated with DpnI, column purified, and cloned into plasmid ML-4059 using Gibson assembly. Gibson products were transformed into DH5α, yielding ∼60,000 colonies that were grown overnight at 37 °C. Overnight culture was prepped to obtain the mutagenized library, which was then electroporated into strain ML-4049 and plated on medium containing 100ng/µL aTc and 100 µM vanillate to induce toxin and *ramF* variants, respectively. The mutagenized library was deep-sequenced pre- and post-selection to identify enriched RamF variants that inhibit MazF at a high induction level. These variants were further validated by constructing new plasmids with single, double or triple mutations on *ramF*.

### Protein structure prediction with AlphaFold2

The predicted structure of the DnaK-RamF complex was generated using AlphaFold2^43, 44^, modeling both proteins as monomers with default parameters (MSA method: mmseqs2, pair mode: unpaired, number of models: 5, max recycles: 3).

## Supplemental Figure Legends

**Fig. S1.**
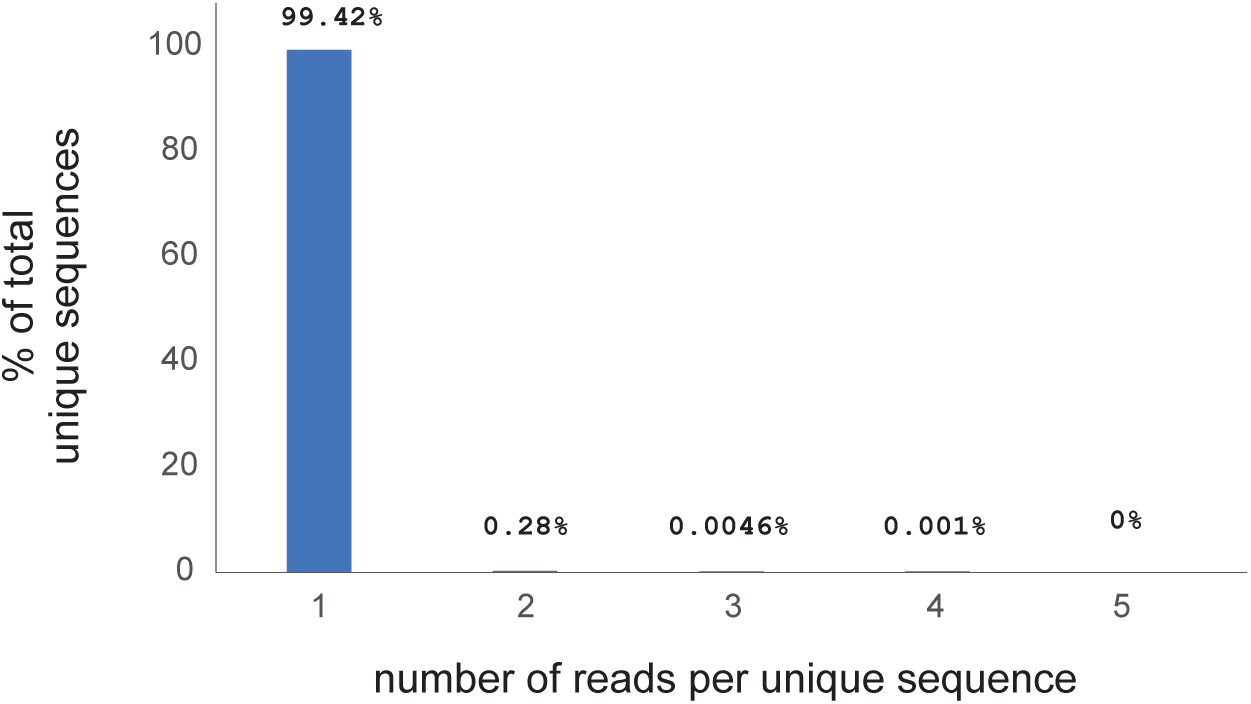
Read counts of the library pre-selection. Number of reads per unique sequence based on deep sequencing of the random protein library pre-selection. The total read count was ∼300,000.

**Fig. S2.**
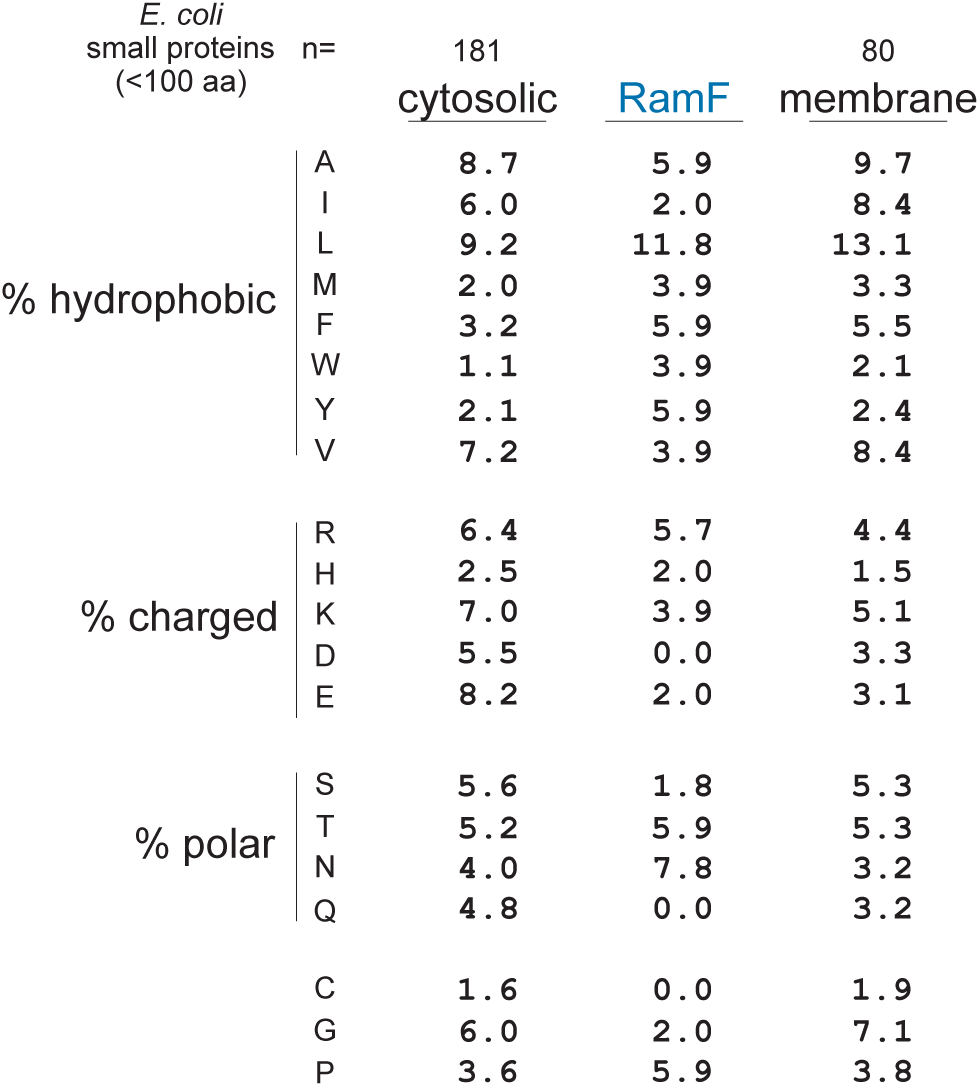
Amino acid composition of RamF. The amino acid composition of RamF compared to cytosolic (n=181) and membrane (n=80) proteins in MG1655 *E. coli* whose lengths are each < 100 amino acids.

**Fig. S3.**
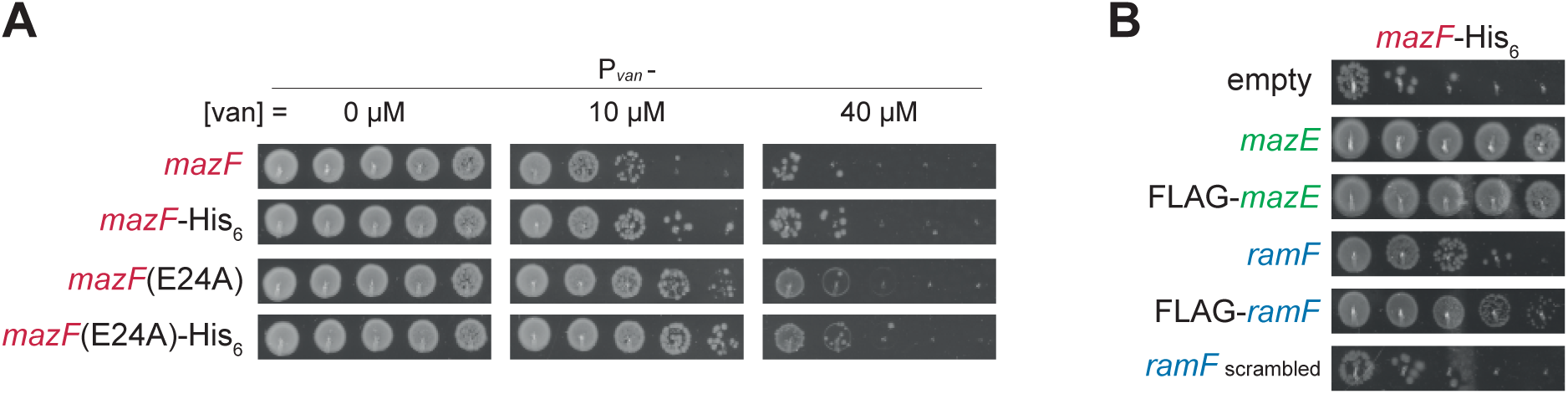
Epitope-tagging of MazF and RamF does not interfere with their functions. (A) 10-fold serial dilution spotting of cells expressing *mazF*, *mazF-His_6_*, *mazF(E24A)*, or *mazF(E24A)-His_6_*, from P*_van_*. (B) 10-fold serial dilution spotting of cells expressing *mazF-His_6_* from P*_van_*. Cells were additionally expressing *mazE*, *FLAG-mazE*, *ramF*, *FLAG-ramF*, scrambled *ramF*, or an empty vector.

**Fig. S4.**
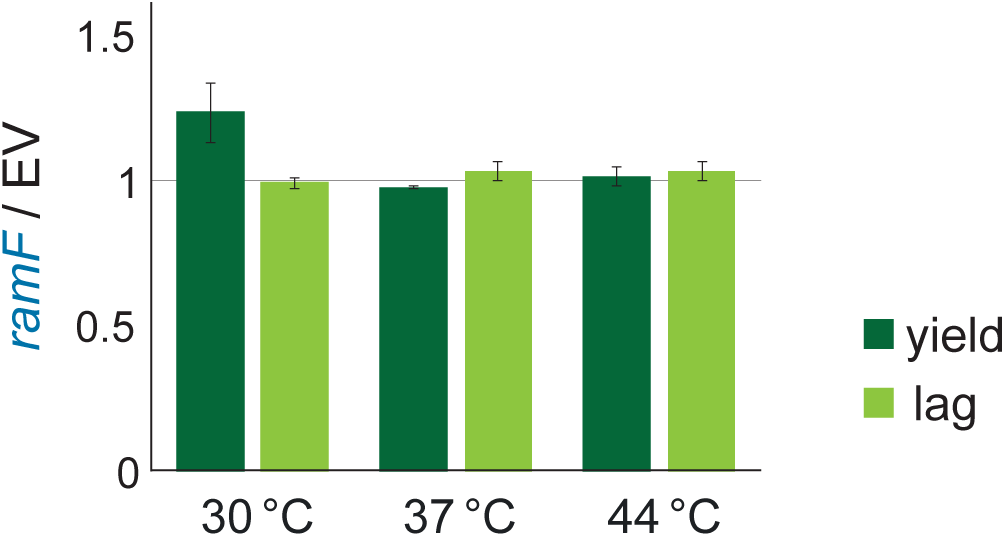
Growth characteristics of cells producing RamF. Lag time (time to reach OD_600_ = 0.2) and culture yield (final OD_600_) ratios between cells producing RamF to empty vector at the growth temperatures indicated.

**Fig. S5.**
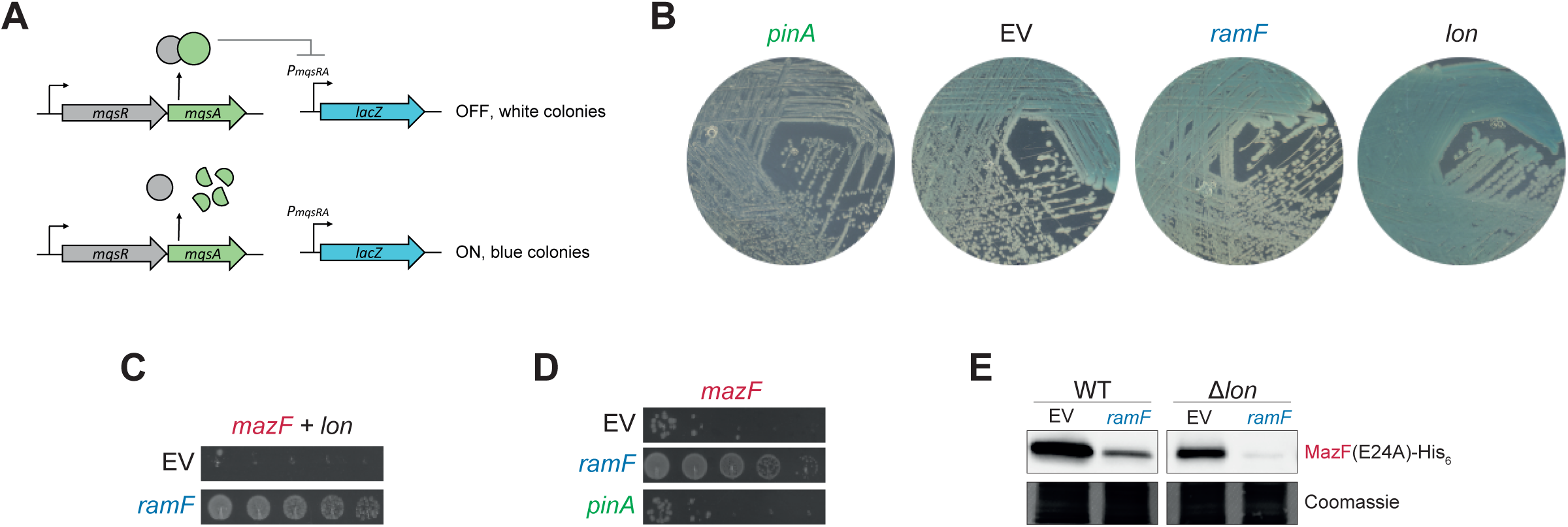
RamF does not inhibit the Lon protease. (A) A system to measure *in vivo* activity of Lon protease: The MqsRA complex inhibits the P*_mqsRA_* promoter driving a *lacZ* reporter. In this system, Lon activity is correlated to LacZ production levels because the antitoxin MqsA is a Lon substrate, and upon antitoxin degradation, LacZ is produced, and colonies turn blue. (B) Cells harboring the system described in (A) also expressing *lon*, *ramF*, *pinA* (a known Lon inhibitor), or an empty vector. (C) 10-fold serial dilution spotting of cells expressing *mazF*, overexpressing *lon*, and additionally expressing *ramF* or an empty vector. (D) 10-fold serial dilution spotting of cells expressing *mazF*, and additionally expressing *ramF*, *pinA*, or an empty vector. (E) Immunoblot of MazF(E24A)-His_6_ expressed from P*_van_*, in control cells or cells lacking the protease Lon. Cells additionally expressing *ramF* or harboring an empty vector. Loading control is based on Coomassie staining of total protein.

**Figure S6.**
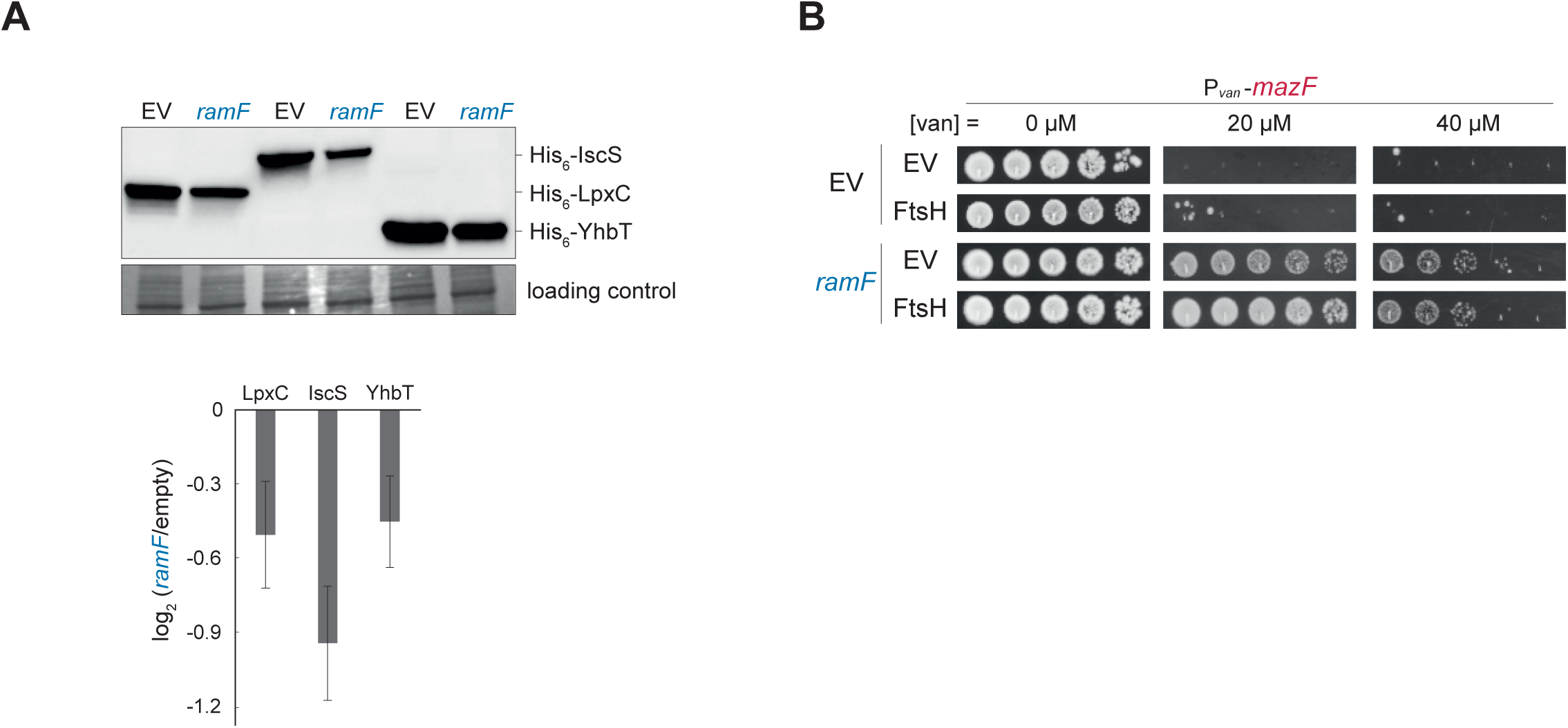
Overproduction of FtsH is insufficient to inhibit MazF and does not alter RamF efficenty as a MazF inhibitor. (A) Immunoblot of His_6_-IscS, His_6_-LpxC, or His_6_-YhbT, known FtsH substrates, from cells co-expressing *ramF* or harboring an empty vector. Loading control is based on Coomassie staining of total protein. (B) 10-fold serial dilution spotting of cells co-expressing (i) *mazF*, (ii) empty vector or *ramF*, and (iii) empty vector or *ftsH*.

**Fig. S7.**
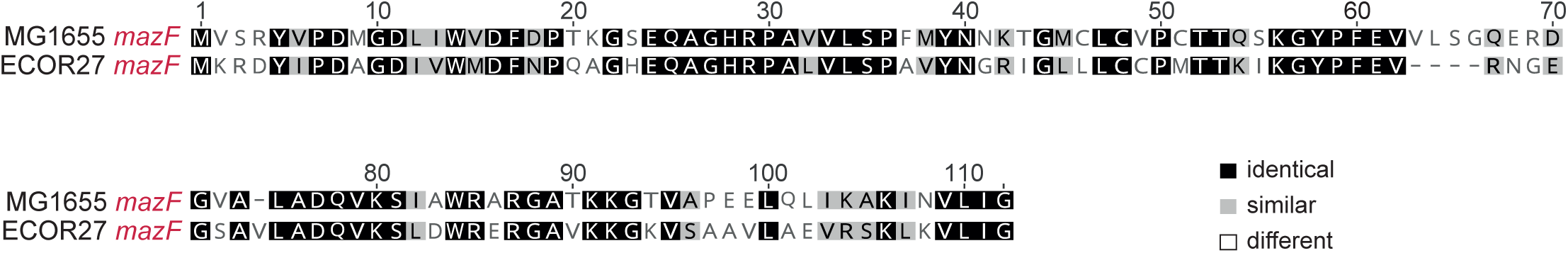
Amino acid alignment of MG1655 MazF and ECOR27 MazF.

## Acknowledgements

We thank the MIT BioMicro Center and its staff for their support in sequencing; the MIT Biopolymers and Proteomics Core and its staff for their help in mass spectrometry experiments; D Ding and C McClune for help with library construction; K. Gozzi, S. Mendoza, A. Murray, S. Srikant, and C. Vassallo for comments on the manuscript; P. DeWeirdt, C. Doering, M. Guzzo, M. LeRoux, T. Zhang, and all members of the Laub laboratory for helpful discussions. IF was supported by a long-term fellowship (LT000706/2018) from the Human Frontier Science Program. MTL is an Investigator of the Howard Hughes Medical Institute.

## References

1. Barrick, J. E. & Lenski, R. E. Genome dynamics during experimental evolution. Nat Rev Genet 14, 827–39 (2013).

2. Tautz, D. & Domazet-Lošo, T. The evolutionary origin of orphan genes. Nat Rev Genet 12, 692– 702 (2011).

3. Andersson, D. I., Jerlström-Hultqvist, J. & Näsvall, J. Evolution of New Functions De Novo and from Preexisting Genes. Cold Spring Harb Perspect Biol 7, a017996 (2015).

4. McLysaght, A. & Hurst, L. D. Open questions in the study of de novo genes: what, how and why. Nat Rev Genet 1–31 (2016) doi:10.1038/nrg.2016.78.

5. Vakirlis, N. et al. De novo emergence of adaptive membrane proteins from thymine-rich genomic sequences. Nat Commun 11, 781 (2020).

6. Vakirlis, N., Carvunis, A.-R. R. & McLysaght, A. Synteny-based analyses indicate that sequence divergence is not the main source of orphan genes. Elife 9, 1–23 (2020).

7. Zhao, L., Saelao, P., Jones, C. D. & Begun, D. J. Origin and spread of de novo genes in Drosophila melanogaster populations. Science 343, 769–72 (2014).

8. Weisman, C. M., Murray, A. W. & Eddy, S. R. Many, but not all, lineage-specific genes can be explained by homology detection failure. PLoS Biol 18, 1–24 (2020).

9. Weisman, C. M., Murray, A. W. & Eddy, S. R. Mixing genome annotation methods in a comparative analysis inflates the apparent number of lineage-specific genes. Curr Biol 32, 2632–2639.e2 (2022).

10. Weisman, C. M. The Origins and Functions of De Novo Genes: Against All Odds? Journal of Molecular Evolution vol. 90 244–257 Preprint at https://doi.org/10.1007/s00239-022-10055-3 (2022).

11. Carvunis, A.-R. et al. Proto-genes and de novo gene birth. Nature 487, 370–4 (2012).

12. Neme, R. & Tautz, D. Fast turnover of genome transcription across evolutionary time exposes entire non-coding DNA to de novo gene emergence. Elife 5, 1–20 (2016).

13. Ruiz-Orera, J., Verdaguer-Grau, P., Villanueva-Cañas, J. L., Messeguer, X. & Albà, M. M. Translation of neutrally evolving peptides provides a basis for de novo gene evolution. Nat Ecol Evol 2, 890–896 (2018).

14. Zhang, L. et al. Rapid evolution of protein diversity by de novo origination in Oryza. Nat Ecol Evol 3, 679–690 (2019).

15. Chen, J. et al. Pervasive functional translation of noncanonical human open reading frames. Science *(1979)* 367, 140–146 (2020).

16. Tretyachenko, V. et al. Random protein sequences can form defined secondary structures and are well-tolerated in vivo. Sci Rep 7, 15449 (2017).

17. Heames, B. et al. Experimental characterisation of de novo proteins and their unevolved random-sequence counterparts. doi:https://doi.org/10.1101/2022.01.14.476368.

18. Wang, M. S. & Hecht, M. H. A Completely de Novo ATPase from Combinatorial Protein Design. J Am Chem Soc 142, 15230–15234 (2020).

19. Wilson, D. S., Keefe, A. D. & Szostak, J. W. The use of mRNA display to select high-affinity protein-binding peptides. Proc Natl Acad Sci U S A 98, 3750–5 (2001).

20. Spangler, L. C. et al. A de novo protein catalyzes the synthesis of semiconductor quantum dots. Proc Natl Acad Sci U S A 119, (2022).

21. Keefe, A. D. & Szostak, J. W. Functional proteins from a random-sequence library. Nature 410, 715–8 (2001).

22. Betanzos, C. M. et al. Bacterial glycoprofiling by using random sequence peptide microarrays. ChemBioChem 10, 877–888 (2009).

23. Knopp, M. et al. De Novo Emergence of Peptides That Confer Antibiotic Resistance. mBio 10, 1–15 (2019).

24. Knopp, M. et al. A novel type of colistin resistance genes selected from random sequence space. PLoS Genet 17, e1009227 (2021).

25. Mutalik, V. K. et al. Precise and reliable gene expression via standard transcription and translation initiation elements. Nat Methods 10, 354–360 (2013).

26. Culviner, P. H. & Laub, M. T. Global Analysis of the E. coli Toxin MazF Reveals Widespread Cleavage of mRNA and the Inhibition of rRNA Maturation and Ribosome Biogenesis. Mol Cell 1–13 (2018) doi:10.1016/j.molcel.2018.04.026.

27. Culviner, P. H., Nocedal, I., Fortune, S. M. & Laub, M. T. Global Analysis of the Specificities and Targets of Endoribonucleases from Escherichia coli Toxin-Antitoxin Systems. mBio e0201221 (2021) doi:10.1128/mBio.02012-21.

28. Brielle, R., Pinel-Marie, M. L. & Felden, B. Linking bacterial type I toxins with their actions. Curr Opin Microbiol 30, 114–121 (2016).

29. Ochman, H. & Selander, R. K. Standard reference strains of Escherichia coli from natural populations. J Bacteriol 157, 690–3 (1984).

30. Zorzini, V. et al. Substrate recognition and activity regulation of the Escherichia coli mRNA endonuclease MazF. Journal of Biological Chemistry 291, 10950–10960 (2016).

31. Li, G. Y. et al. Characterization of dual substrate binding sites in the homodimeric structure of Escherichia coli mRNA interferase MazF. J Mol Biol 357, 139–150 (2006).

32. Kamada, K., Hanaoka, F. & Burley, S. K. Crystal structure of the MazE/MazF complex: Molecular bases of antidote-toxin recognition. Mol Cell 11, 875–884 (2003).

33. Ogura, T. et al. Balanced biosynthesis of major membrane components through regulated degradation of the committed enzyme of lipid A biosynthesis by the AAA protease FtsH (HflB) in Escherichia coli. Mol Microbiol 31, 833–844 (1999).

34. Zorzini, V. et al. Escherichia coli antitoxin MazE as transcription factor: Insights into MazE-DNA binding. Nucleic Acids Res 43, 1241–1256 (2015).

35. Schramm, F. D., Schroeder, K. & Jonas, K. Protein aggregation in bacteria. FEMS Microbiol Rev 44, 54–72 (2019).

36. Tomoyasu, T., Mogk, A., Langen, H., Goloubinoff, P. & Bukau, B. Genetic dissection of the roles of chaperones and proteases in protein folding and degradation in the Escherichia coli cytosol. Mol Microbiol 40, 397–413 (2001).

37. Mogk, A., Deuerling, E., Vorderwülbecke, S., Vierling, E. & Bukau, B. Small heat shock proteins, ClpB and the DnaK system form a functional triade in reversing protein aggregation. Mol Microbiol 50, 585–595 (2003).

38. Chapman, E. et al. Global aggregation of newly translated proteins in an Escherichia coli strain deficient of the chaperonin GroEL. Proc Natl Acad Sci U S A 103, 15800–15805 (2006).

39. Zhao, K., Liu, M. & Burgess, R. R. The global transcriptional response of Escherichia coli to induced σ32 protein involves σ32 regulon activation followed by inactivation and degradation of σ32 in vivo. Journal of Biological Chemistry 280, 17758–17768 (2005).

40. Patra, M., Roy, S. S., Dasgupta, R. & Basu, T. GroEL to DnaK chaperone network behind the stability modulation of σ32 at physiological temperature in Escherichia coli. FEBS Lett 589, 4047–4052 (2015).

41. Nonaka, G., Blankschien, M., Herman, C., Gross, C. A. & Rhodius, V. A. Regulon and promoter analysis of the E. coli heat-shock factor, σ32, reveals a multifaceted cellular response to heat stress. Genes Dev 20, 1776–1789 (2006).

42. Yura, T. et al. Analysis of σ32 mutants defective in chaperone-mediated feedback control reveals unexpected complexity of the heat shock response. Proc Natl Acad Sci U S A 104, 17638–17643 (2007).

43. Mirdita, M. et al. ColabFold: making protein folding accessible to all. Nat Methods 19, 679–682 (2022).

44. Jumper, J. et al. Highly accurate protein structure prediction with AlphaFold. Nature 596, 583– 589 (2021).

45. Mayer, M. P., Rudiger, S. & Bukau, B. Molecular basis for interactions of the DnaK chaperone with substrates. Biol Chem 381, 877–885 (2000).

46. Zhu, X. et al. Structural analysis of substrate binding by the molecular chaperone DnaK. Science 272, 1606–14 (1996).

47. Pellecchia, M. et al. Structural insights into substrate binding by the molecular chaperone DnaK. Nat Struct Biol 7, 298–303 (2000).

48. Bittner, L. M., Arends, J. & Narberhaus, F. When, how and why? Regulated proteolysis by the essential FtsH protease in Escherichia coli. Biol Chem 398, 625–635 (2017).

49. Führer, F., Langklotz, S. & Narberhaus, F. The C-terminal end of LpxC is required for degradation by the FtsH protease. Mol Microbiol 59, 1025–1036 (2006).

50. Herman, C., Thévenet, D., Bouloc, P., Walker, G. C. & D’Ari, R. Degradation of carboxy-terminal-tagged cytoplasmic proteins by the Escherichia coli protease HflB (FtsH). Genes Dev 12, 1348–1355 (1998).

51. Bittner, L. M., Westphal, K. & Narberhaus, F. Conditional proteolysis of the membrane protein YfgM by the FtsH protease depends on a novel N-terminal degron. Journal of Biological Chemistry 290, 19367–19378 (2015).

52. Ruiz-Orera, J. & Albà, M. M. Translation of Small Open Reading Frames: Roles in Regulation and Evolutionary Innovation. Trends in Genetics 35, 186–198 (2019).

53. Couso, J. P. & Patraquim, P. Classification and function of small open reading frames. Nat Rev Mol Cell Biol 18, 575–589 (2017).

54. Storz, G., Wolf, Y. I. & Ramamurthi, K. S. Small proteins can no longer be ignored. Annu Rev Biochem 83, 753–777 (2014).

55. Orr, M. W., Mao, Y., Storz, G. & Qian, S. B. Alternative ORFs and small ORFs: shedding light on the dark proteome. Nucleic Acids Res 48, 1029–1042 (2020).

56. Weaver, J., Mohammad, F., Buskirk, A. R. & Storz, G. Identifying Small Proteins by Ribosome Profiling with Stalled Initiation Complexes. mBio 10, (2019).

57. Stringer, A., Smith, C., Mangano, K. & Wade, J. T. Identification of Novel Translated Small Open Reading Frames in Escherichia coli Using Complementary Ribosome Profiling Approaches. J Bacteriol 204, (2022).

58. Sberro, H. et al. Large-Scale Analyses of Human Microbiomes Reveal Thousands of Small, Novel Genes. Cell 178, 1245–1259.e14 (2019).

59. Miravet-Verde, S., et al. Unraveling the hidden universe of small proteins in bacterial genomes. Mol Syst Biol 15, 1–17 (2019).

60. Kemp, G. & Cymer, F. Small membrane proteins - Elucidating the function of the needle in the haystack. Biol Chem 395, 1365–1377 (2014).

61. Guerzoni, D. & McLysaght, A. De novo genes arise at a slow but steady rate along the primate lineage and have been subject to incomplete lineage sorting. Genome Biol Evol 8, 1222–1232 (2016).

62. Vakirlis, N., Vance, Z., Duggan, K. M. & McLysaght, A. De novo birth of functional microproteins in the human lineage. Cell Rep 41, (2022).

63. Neme, R., Amador, C., Yildirim, B., McConnell, E. & Tautz, D. Random sequences are an abundant source of bioactive RNAs or peptides. Nat Ecol Evol 1, 0217 (2017).

64. Weisman, C. M. & Eddy, S. R. Gene Evolution: Getting Something from Nothing. Curr Biol 27, R661–R663 (2017).

65. Knopp, M. & Andersson, D. I. No beneficial fitness effects of random peptides. Nat Ecol Evol 2, 1046–1047 (2018).

66. Digianantonio, K. M. & Hecht, M. H. A protein constructed de novo enables cell growth by altering gene regulation. Proc Natl Acad Sci U S A 113, 2400–5 (2016).

67. Hoegler, K. J. & Hecht, M. H. A de novo protein confers copper resistance in Escherichia coli. Protein Science 1249–1259 (2016) doi:10.1002/pro.2871.

68. Jurėnas, D., Fraikin, N., Goormaghtigh, F. & van Melderen, L. Biology and evolution of bacterial toxin–antitoxin systems. Nat Rev Microbiol 0123456789, (2022).

69. Burga, A., Ben-David, E. & Kruglyak, L. Toxin-Antidote Elements Across the Tree of Life. Annu Rev Genet 54, 1–29 (2020).

70. Loris, R. & Garcia-Pino, A. Disorder-and dynamics-based regulatory mechanisms in toxin-antitoxin modules. Chem Rev 114, 6933–6947 (2014).

71. Garcia-Pino, A. et al. Doc of prophage P1 is inhibited by its antitoxin partner Phd through fold complementation. J Biol Chem 283, 30821–7 (2008).

72. de Gieter, S. et al. The intrinsically disordered domain of the antitoxin Phd chaperones the toxin Doc against irreversible inactivation and misfolding. J Biol Chem 289, 34013–23 (2014).

73. Cherny, I. & Gazit, E. The YefM antitoxin defines a family of natively unfolded proteins: implications as a novel antibacterial target. J Biol Chem 279, 8252–61 (2004).

74. Snead, K. J., Moore, L. L. & Bourne, C. R. ParD Antitoxin Hotspot Alters a Disorder-to-Order Transition upon Binding to Its Cognate ParE Toxin, Lessening Its Interaction Affinity and Increasing Its Protease Degradation Kinetics. Biochemistry 61, 34–45 (2022).

75. Wilson, B. A., Foy, S. G., Neme, R. & Masel, J. Young genes are highly disordered as predicted by the preadaptation hypothesis of de novo gene birth. Nat Ecol Evol 1, (2017).

76. Kosinski, L. J., Aviles, N. R., Gomez, K. & Masel, J. Random Peptides Rich in Small and Disorder-Promoting Amino Acids Are Less Likely to Be Harmful. Genome Biol Evol 14, (2022).

77. Cui, Y. et al. Bacterial MazF/MazE toxin-antitoxin suppresses lytic propagation of arbitrium-containing phages. Cell Rep 41, 111752 (2022).

78. LeRoux, M. et al. The DarTG toxin-antitoxin system provides phage defence by ADP-ribosylating viral DNA. Nat Microbiol (2022) doi:10.1038/s41564-022-01153-5.

79. LeRoux, M. & Laub, M. T. Toxin-Antitoxin Systems as Phage Defense Elements. Annu Rev Microbiol 76, 21–43 (2022).

80. Zhang, T. et al. Direct activation of a bacterial innate immune system by a viral capsid protein. Nature (2022) doi:10.1038/s41586-022-05444-z.

81. Short, F. L., Akusobi, C., Broadhurst, W. R. & Salmond, G. P. C. The bacterial Type III toxin-antitoxin system, ToxIN, is a dynamic protein-RNA complex with stability-dependent antiviral abortive infection activity. Sci Rep 8, (2018).

82. Otsuka, Y. & Yonesaki, T. Dmd of bacteriophage T4 functions as an antitoxin against Escherichia coli LsoA and RnlA toxins. Mol Microbiol 83, 669–681 (2012).

83. Srikant, S., Guegler, C. K. & Laub, M. T. The evolution of a counter-defense mechanism in a virus constrains its host range. Elife 11, (2022).

84. Fremin, B. J. et al. Thousands of small, novel genes predicted in global phage genomes. Cell Rep 39, 110984 (2022).

85. Luo, Y., Zhang, T. & Wu, H. The transport and mediation mechanisms of the common sugars in Escherichia coli. Biotechnol Adv 32, 905–19 (2014).

86. Schleif, R. AraC protein, regulation of the l-arabinose operon in Escherichia coli, and the light switch mechanism of AraC action. FEMS Microbiol Rev 34, 779–96 (2010).

87. Schleif, R. Regulation of the L-arabinose operon of Escherichia coli. Trends Genet 16, 559–65 (2000).

88. Khlebnikov, A., Datsenko, K. A., Skaug, T., Wanner, B. L. & Keasling, J. D. Homogeneous expression of the P(BAD) promoter in Escherichia coli by constitutive expression of the low-affinity high-capacity AraE transporter. Microbiology (Reading) 147, 3241–7 (2001).

89. Datsenko, K. a & Wanner, B. L. One-step inactivation of chromosomal genes in Escherichia coli K-12 using PCR products. Proc Natl Acad Sci U S A 97, 6640–5 (2000).

90. Welinder, C. & Ekblad, L. Coomassie staining as loading control in Western blot analysis. J Proteome Res 10, 1416–1419 (2011).

91. Culviner, P. H., Guegler, C. K. & Laub, M. T. A Simple, Cost-Effective, and Robust Method for rRNA Depletion in RNA-Sequencing Studies. mBio 11, (2020).

92. Dillies, M. A. et al. A comprehensive evaluation of normalization methods for Illumina high-throughput RNA sequencing data analysis. Brief Bioinform 14, 671–683 (2013).

93. Ducret, A., Quardokus, E. M. & Brun, Y. V. MicrobeJ, a tool for high throughput bacterial cell detection and quantitative analysis. Nat Microbiol 1, (2016).

94. Srikant, S., Gaudet, R. & Murray, A. W. Selecting for Altered Substrate Specificity Reveals the Evolutionary Flexibility of ATP-Binding Cassette Transporters. Current Biology 30, 1689–1702.e6 (2020).

